# A recently evolved TAF8 isoform arising from an *Alu* insertion increases TFIID assembly complexity in the human lineage

**DOI:** 10.64898/2026.01.19.699933

**Authors:** Andrea Bernardini, Alberto Gallo, Elizabeth Scheer, Bastien Morlet, Diletta Dolfini, Roberto Mantovani, Stéphane D. Vincent, László Tora

## Abstract

Despite its centrality in regulating RNA polymerase II transcription in all eukaryotes, the TFIID general transcription factor exhibits several layers of variability across different tissues and developmental stages, representing an underexplored hub of evolutionary innovation. Here, we describe a novel short isoform of TAF8 (TAF8s) – a TFIID scaffold subunit – which evolved from the use of an intronic polyadenylation site (iPAS) in the human lineage. Comparative genomics analyses show that the emergence of *TAF8s* expression in the human lineage coincides with minute DNA changes in the iPAS at the edge of a FLAM-C *Alu* element that inserted into intron 5 in the common ancestor of anthropoid primates (Simiiformes). The human-specific *TAF8s* isoform lacks nearly half of the canonical coding sequence, is widely expressed across human tissues, and constitutes a substantial fraction of the *TAF8* transcript pool. *TAF8s* is translated into a truncated protein isoform that lacks the nuclear-localization signal and localises in the cytoplasm. TAF8s interacts with its histone-fold partner TAF10 and other core TFIID subunits, while entirely losing its interactions with TAF2, thus giving rise to alternative TFIID sub-complexes in human cells. Our study suggests that the TFIID complex underwent a recent diversification through a stepwise evolutionary acquisition of complexity in the *TAF8* locus in the human lineage, leading to the expression of a novel truncated pan-isoform that might work as a modulator of TFIID assembly. We discuss the ramifications of our findings in TFIID complex diversification, its evolvability, and the genetics of TAF-related congenital disorders.

## Introduction

Evolutionary changes in proteins involved in transcriptional regulation can trigger broad effects on the activity of many genes, thus representing key molecular ‘knobs’ that can quickly shape evolutionary trajectories (Johnson 2017). The basal transcription machinery – including the General Transcription Factors (GTFs) and RNA polymerase II (Pol II) – assembles at core promoters into a pre-initiation complex (PIC) that integrates inputs from multiple molecular sources to kickstart transcription. Despite the ‘general’ appellative, the GTF composition can vary depending on developmental stages and the tissues considered, especially through paralog-switching mechanisms (Levine and Tjian 2003). The 1.2 MDa TFIID complex is the GTF that recognises core promoter elements, nucleates PIC assembly, and regulates its activity (Verrijzer et al. 1995; Malik and Roeder 2023). In metazoa, TFIID is composed of the TATA-binding protein (TBP) and 13 TBP-associated factors (TAFs) (Tora 2002). The complex is constituted of three structural lobes (A, B, C) and contains a set of five TAFs present in double copy (TAF4, TAF5, TAF6, TAF9, TAF12), named core TFIID. TAF8 works as a scaffold protein that physically connects TFIID lobe B to lobe C (Patel et al. 2018; Chen et al. 2021). TFIID complex assembly takes place in the cytoplasm through a complex series of hierarchical steps, many of which occur during the translation of key subunits (Kamenova et al. 2019; Bernardini et al. 2023; Bernardini and Tora 2024).

All the tested *TAFs* (*Taf1*, *Taf4*, *Taf5*, *Taf7*, *Taf8*, *Taf10, Taf12, Taf13*) are essential genes, as mouse knockouts (KO) die at the embryonic stage, mostly around peri-implantation (Voss et al. 2000; Mohan et al. 2003; Gegonne et al. 2012; Langer et al. 2016; Crombie, Korecki, et al. 2024; He et al. 2024; Martianov et al. 2025). The essentiality of *TAF*s has also been shown in zebrafish (*taf1*, *taf1b*, *taf2*, *taf5*, *taf7*, *taf8*) and budding yeast (Shen et al. 2003; Amsterdam et al. 2004; Gudmundsson et al. 2019; Leid et al. 2023).

The origin of TFIID predates the last eukaryotic common ancestor (LECA), and although the minimal domain architecture of its subunits appears rather conserved, it also displays a high degree of plasticity in the length and sequence of the interdomain regions (Antonova et al. 2019). Despite its foundational role in PIC assembly, TFIID is a hub of evolutionary innovation in metazoan evolution. Several of its subunits, including TBP, underwent gene duplications (Akhtar and Veenstra 2011; Martianov et al. 2016; Yu et al. 2020). The resulting paralog genes followed either partial functional overlap (co-expression with mostly conservative functions) or sub-functionalization with tissue-specific expression, especially in gonads (Müller and Tora 2004; Antonova et al. 2019; Gura et al. 2020). The existence of these paralog genes and corresponding specialized TFIID variants suggests that in metazoans TFIID is not a monolithic entity and PIC assembly is a multifaceted process (Bell and Tora 1999; Müller and Tora 2004; Levine et al. 2014). In this regard, a second source of variability of the basal transcription machinery came from the detection of partial TFIID complexes in cellular extracts, where one subunit or entire sub-modules are missing or found at sub-stoichiometric levels, although it remains difficult to discern whether they are active functional units or mere assembly intermediates (Brou, Wu, et al. 1993; Brou, Chaudhary, et al. 1993; Brou, Wu, et al. 1993; Jacq et al. 1994; Martin et al. 1999; Bell et al. 2001; Hardy et al. 2002). In this regard, upon experimental depletion of TAF7 or TAF10 in mouse embryonic stem cells (mESCs), the remaining partial complexes seem to be able to support TBP recruitment and nascent transcription for several cell divisions, yet they do not sustain mESCs viability in the long term, with *Taf10* KO being more severe than *Taf7* KO (Hisler et al. 2024). Another study showed that *Taf13* KO mESCs were viable, with TBP recruitment and gene expression only mildly affected, despite showing reduced Pol II occupancy, growth defects, and blocked differentiation potential (Martianov et al. 2025). Indeed, there is evidence that not all TFIID subunits contribute equally to transcription (Sun et al. 2021).

Strikingly, there is evidence that *TAF*s tested with conditional mouse knockouts (*Taf1*, *Taf7*, *Taf10*) become dispensable in differentiated adult tissues (skin, liver, thymus, hematopoietic system), while being essential during the development of those same tissues (Indra et al. 2005; Tatarakis et al. 2008; Gegonne et al. 2012; Liu et al. 2025). An additional clue of the differential *in vivo* requirements of *TAF*s came from the discovery of rare congenital disorders caused by pathogenic variants in several *TAF* genes, including *TAF1*, *TAF2*, *TAF4*, *TAF6*, *TAF8,* and *TAF13*, and characterized by neurological disorders (Hellman-Aharony et al. 2013; Alazami et al. 2015; Tawamie et al. 2017; El-Saafin et al. 2018; Janssen et al. 2022; Crombie, Cleverley, et al. 2024). The existence of these patients demonstrates that none of those *TAF* variants are embryonic lethal, yet the main process disrupted by these disorders – now collectively called TAFopathies – is brain development, suggesting that embryonic and adult tissues are differentially susceptible to defects in TFIID activity.

The expression of different *TAF* isoforms through alternative promoters, splicing, or poly-adenylation sites represents another vast – still underexplored – source of plasticity for TFIID evolutionary innovations. Here, we describe the existence of a novel, truncated TAF8 protein isoform that recently evolved in the human lineage. We characterize the origin, expression, evolutionary track, subcellular localization, structural features, and interacting subunits of this novel isoform. We discuss the ramifications of our discovery on TFIID complex diversification and the interpretation of *TAF8*-related genetic disorders.

## Results

### The human *TAF8* gene produces an abundant uncharacterized short *TAF8* pan-isoform

The number of alternative protein-coding isoforms across TFIID subunit genes appears to be rather heterogeneous, ranging from genes with no alternative products (e.g., *TAF3*, *TAF7*, *TAF9*) to those producing several annotated alternative isoforms (e.g., *TAF1*, *TAF2*, *TAF6*, *TAF8*, *TAF12*). *TAF8* caught our attention, since amongst the several annotated protein-coding transcripts (**Fig. S1A**), two products are widely co-expressed in human tissues (**Fig. 1A**). The first transcript corresponds to the widely studied *TAF8* canonical isoform (*TAF8c*), the second to an uncharacterized shorter transcript, which we name here *TAF8 short* (*TAF8s*). The *TAF8* canonical transcript is constituted of 9 exons in total, including the terminal exon containing a long 3’ untranslated region (UTR) (**Fig. S1A**), encoding the canonical TAF8 protein (310 aa). Instead, *TAF8s* only includes exons 1 to 5, and it is predicted to code for a dramatically shorter protein product (174 aa).

**Figure 1.**
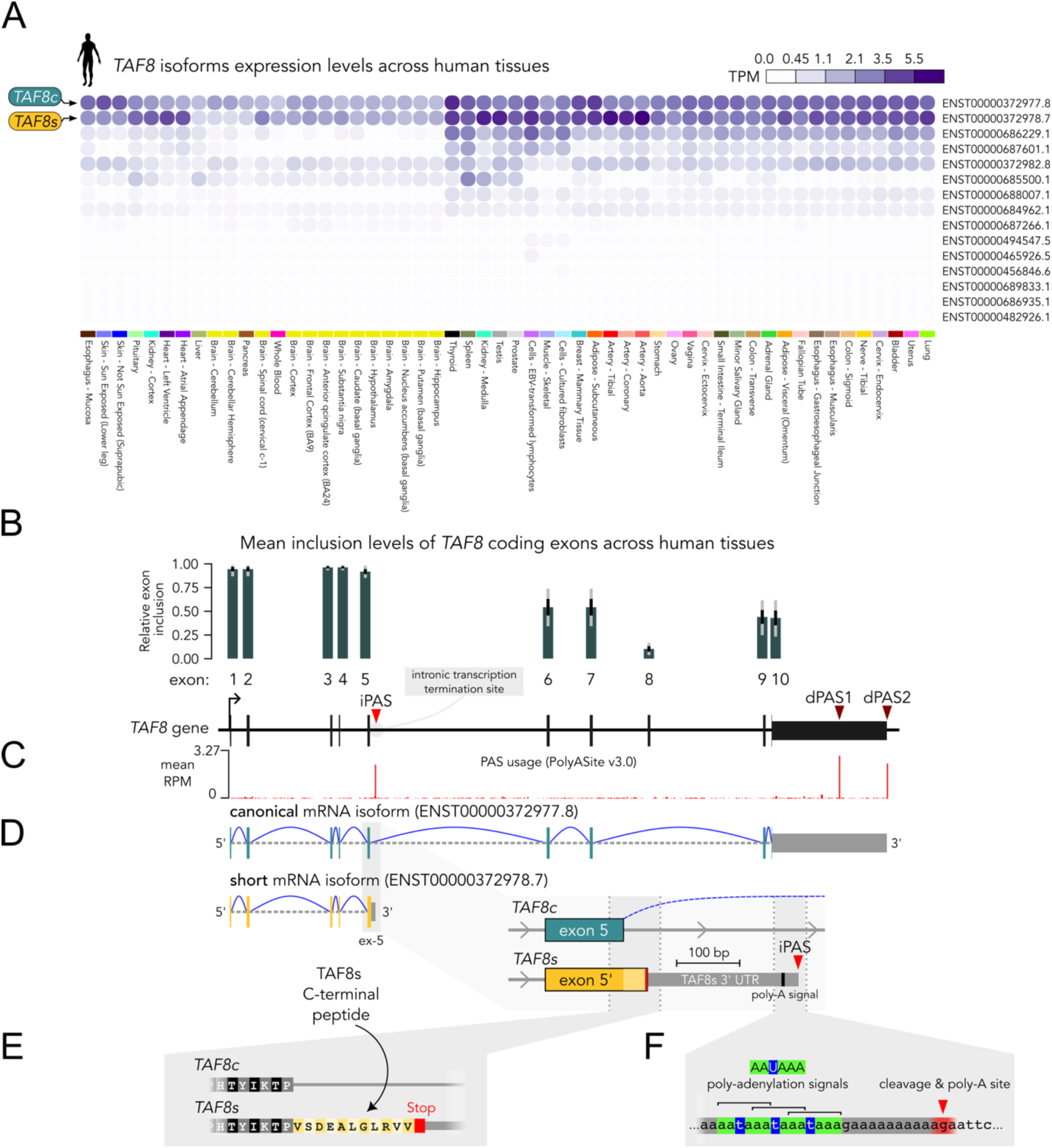
The human *TAF8* locus produces a truncated short isoform using an intronic polyadenylation site (iPAS). **A**. Human *TAF8* isoform gene expression profiles for adult tissues from the GTEx project. *TAF8* canonical and short isoforms are indicated. TPM: transcripts per million. **B**. Human *TAF8* gene exon-intron organization. Mean relative exon inclusion fraction across human tissues is displayed on top of each exon. The data correspond to the pext (proportion expressed across transcripts) score from gnomAD. Error bars represent standard deviation (black) and data range (gray). **C**. PAS usage (3’-seq signal) along *TAF8* gene from PolyASite database (v3.0). The corresponding PASs are indicated above with red arrowheads. **D**. Anatomy of *TAF8* canonical and short isoforms. The inset shows the 3’UTR specific to *TAF8s* resulting from the usage of the iPAS. **E**. Predicted peptide sequence of the C-terminal tail unique to the TAF8s isoform. **F**. Expanded view of the iPAS preceded by a set of three partially overlapping poly-adenylation signals (AAUAAA).

Overall, the expression levels of *TAF8c* and *TAF8s* mRNAs are of the same order of magnitude and together account for the majority of the gene output, whereas the remaining annotated isoforms are only marginally expressed (**Fig. 1A**). The expression levels of the canonical and short isoforms vary slightly depending on the tissue (**Fig. S1B**), with isoforms ratios restricted in the 0.5-2-fold range across most tissues (**Fig. S1C**). The levels of the short isoform can even exceed those of the canonical one, as in heart and testis. Overall, the ubiquitous expression pattern of *TAF8s* makes this uncharacterized gene product an actual *TAF8* pan-isoform.

### An intronic polyadenylation event generates the *TAF8* short isoform

Based on the above observations, we hypothesized that the *TAF8s* isoform originates from a premature transcription termination event, occurring within intron 5 of the *TAF8* gene.

Indeed, the inclusion levels of *TAF8* coding exons across human tissues abruptly decrease by ∼50% downstream of exon 5, on average (**Fig. 1B**). This suggests that a significant proportion of the transcripts are terminated within intron 5, due to an intronic polyadenylation event. The inspection of 3’-sequencing tracks from the PolyASite database confirms the presence of an intronic polyadenylation site (iPAS) ∼280 bp downstream of exon 5 (**Fig. 1C**), in addition to the two major distal sites at the end of the gene (dPAS1-2). Transcript polyadenylation at the iPAS followed by transcription termination generates the *TAF8s* transcript, including a specific ∼240 nt 3’UTR corresponding to the initial region of canonical intron 5 (**Fig. 1D**). Translation of *TAF8s* transcript would result in a truncated TAF8 protein, identical to canonical TAF8 up to positions encoded by exon 5, but lacking the entire C-terminal half of the protein. Importantly, *TAF8s* CDS extends into intron 5, up to an in-frame Stop codon, generating a short C-terminal peptide (11 residues) unique to TAF8s (**Fig. 1E**).

Further downstream in intron 5, we found a series of three – partially overlapping – canonical polyadenylation signals (AATAAA) (**Fig. 1F**). As expected, the predicted cleavage and polyadenylation site is located ∼20 nt downstream of the first AATAAA hexamer (**Fig. S2**).

Moreover, we retrieved EST clones that contained portions of the actual poly-A tails connected to the predicted PAS (**Fig. S2**, bottom panel alignment), substantiating the usage of this intronic polyadenylation site. In summary, *TAF8s* isoform is produced from transcription that ends prematurely within intron 5 due to a series of three polyadenylation signals that trigger nascent transcript cleavage and polyadenylation.

### TAF8s protein is expressed in human cells

Next, we inspected ribosome footprinting (Ribo-seq) datasets and found that *TAF8s* could be translated. Focusing on *TAF8* exon 5, it is evident that the short intronic region corresponding to the 11-residue tail unique to the short isoform is covered by the ribosome protection signal, which ends in correspondence to the Stop codon (**Fig. 2A**).

**Figure 2.**
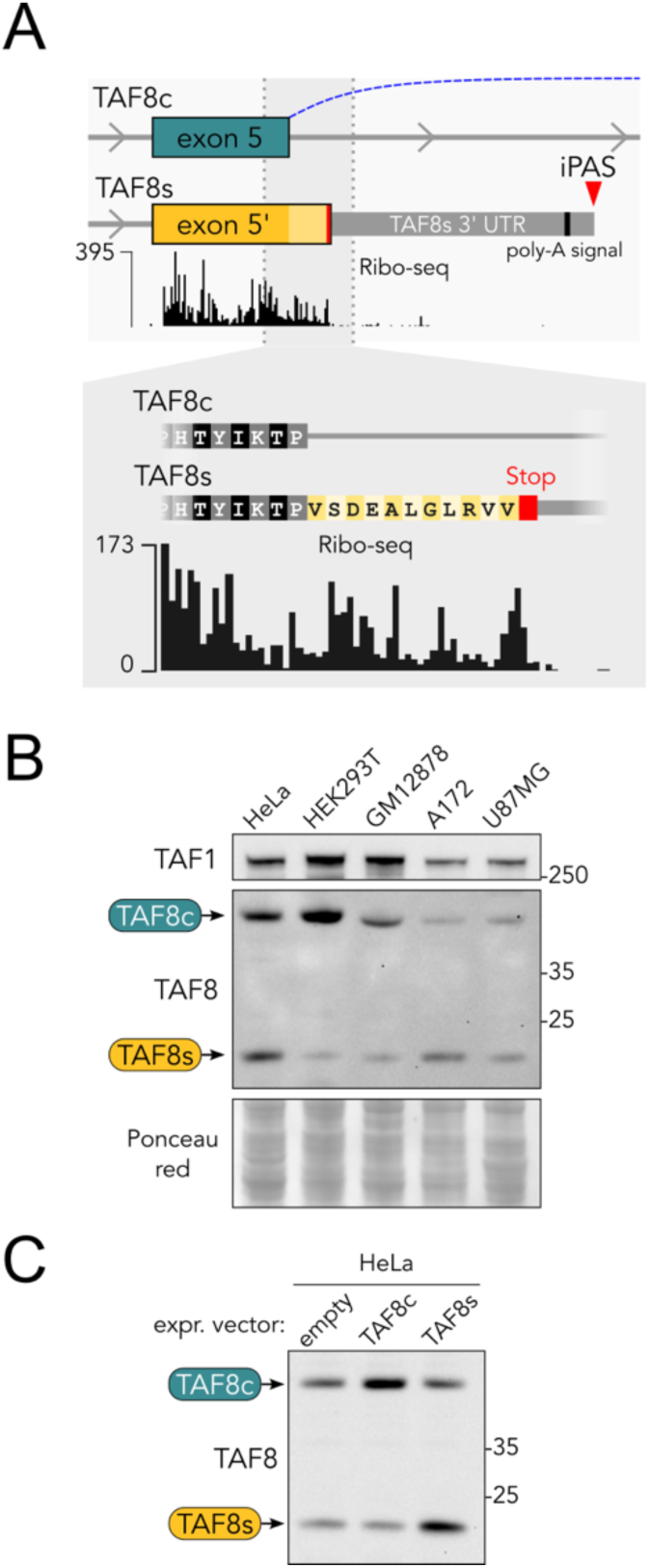
TAF8s is expressed at the protein level. **A**. Aggregate ribosome footprinting (Ribo-seq) signal track focused on *TAF8* exon 5 from the GWIPS-viz browser. The enlarged inset shows active translation of the TAF8s specific tail. **B**. Western blot on RIPA extracts from different human cell lines using an antibody able to recognize both TAF8 isoforms. **C**. TAF8 western blot on RIPA extracts from HeLa cells transfected with TAF8c or TAF8s expression vectors. Empty vector was used as control.

To directly detect the existence of the TAF8s protein, we selected a commercial antibody raised against the TAF8 N-terminal region, common to the TAF8c and TAF8s isoforms. We performed western blot analyses on several human cell lines of different origins using the above antibody. Indeed, apart from the usual band corresponding to TAF8c, we detected a lower band migrating at the predicted molecular weight for TAF8s (∼19 kDa, **Fig. 2B**).

To further test the identity of the protein species detected, we cloned and expressed the CDS of *TAF8c* or *TAF8s* in HeLa cells by transient transfection. As expected, the bands of the exogenously expressed isoforms coincide with the endogenous bands detected in the control sample (“empty” in **Fig. 2C**). We conclude that *TAF8s* is expressed also at the protein level in human cells. Moreover, it represents a relevant proportion of the total TAF8 abundance, opening questions about the molecular features, interactions, conservation, and evolutionary origin of TAF8s.

### TAF8s is a human-specific TAF8 isoform

TAF8 is a highly conserved protein, with a mouse-human sequence identity of 94%. Thus, we investigated whether *TAF8s* isoform expression is also conserved in other species. *TAF8s* isoform is automatically annotated only in human (ENSEMBL transcript ID ENST00000372978), but not in mouse or other vertebrate species. Inspection of uniformly processed RNA-seq tracks from human, rhesus macaque (*Macaca mulatta*) and mouse tissues (Li et al. 2018) shows the expected reduction in exon coverage downstream of exon 5 in human samples, due to premature transcription termination and *TAF8s* production, but not in rhesus or mouse samples (**Fig. 3A**). Accordingly, no signs of exon 5 read-through into the adjacent intron are visible, except from the human samples, where the signal corresponding to *TAF8s* 3’UTR in intron 5 is clearly present (**Fig. 3B**).

**Figure 3.**
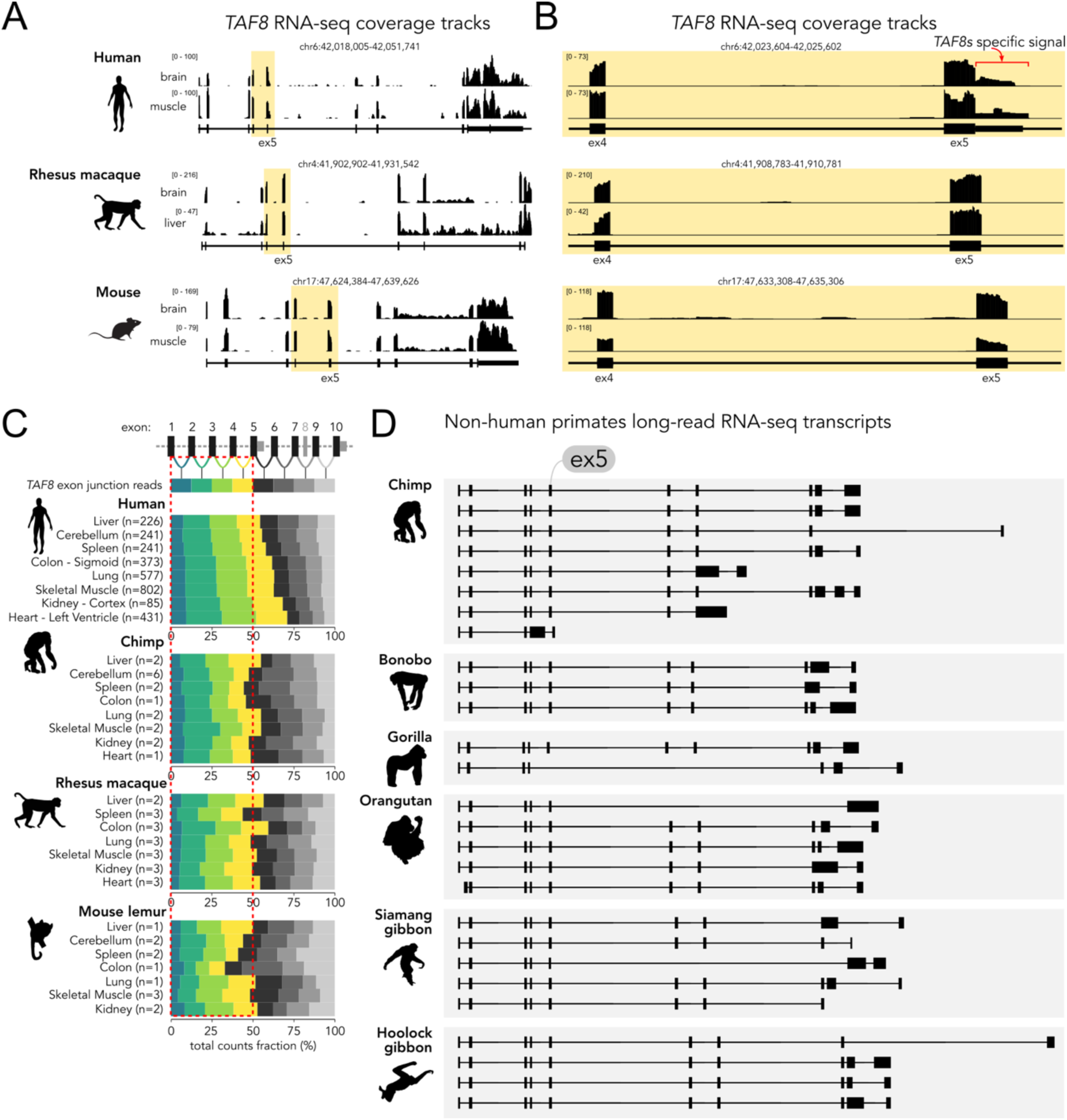
*TAF8* expression data from human and other species. **A**. RNA-seq coverage tracks from human, rhesus macaque, and mouse *TAF8 loci* from (Li et al. 2018). The positions enclosing exons 5 and 6 are highlighted in yellow. **B**. Enlarged view of the regions highlighted in panel A. Reads mapping to *TAF8s* 3’UTR are indicated in red. **C**. Percentages of *TAF8* RNA-seq reads mapping to different exon-exon junctions from multiple primate species and tissues. Exon junctions not present in the *TAF8s* isoform are shaded in grey. A red dashed box indicates the expected proportion of junction reads for exons which are common between *TAF8c* and *TAF8s* (exons 1-5), assuming equal reads distribution along the gene as indicated in the top row. **D**. Exon-intron structures of individual long reads mapped on the *TAF8* gene from different primate species from long-read RNA-seq data.

We further analysed other primate RNA-seq datasets from different tissues and quantified the number of exon-exon junction spanning reads across the *TAF8* gene (**Fig. 3C**). If a substantial proportion of *TAF8s* is produced, one expects to find more reads mapping to exons 1-5 than to the downstream exons 6-10, as the latter are not part of the short isoform. Indeed, this is the case in human samples (**Fig. 3C**), where a clear bias towards the upstream exons 1-5 is visible in several tissues, especially in heart and kidney, where the short isoform is predominant (see also **Fig. S1B**). Instead, in the other primate species analysed – chimp, rhesus macaque, and mouse lemur – we do not find a clear bias towards the upstream exons (**Fig. 3C**), suggesting no substantial expression of *TAF8s*. The same trend can be appreciated by direct inspection of the RNA-seq coverage tracks (**Fig. S3**), where the lower exon coverage downstream of exon 5 and the exon 5 bleed-through reads in intron 5 are not found in non-human primate samples, including the closely related chimpanzee samples. To assess the existence of *TAF8s* isoform more directly, we uniformly analysed available long-read RNA-seq datasets from human and several non-human primate species (chimp, bonobo, gorilla, orangutan, siamang gibbon, and hoolock gibbon). While we detected several instances of the *TAF8s* isoform in human samples (**Fig. S4**), no long reads corresponding to *TAF8s* could be found in non-human primate samples (**Fig. 3D**). Along the same line, we found no evidence of *TAF8s* expression in the African green monkey cell line (COS-7), neither at mRNA (**Fig. S5A**) nor at protein level (**Fig. S5B**). Overall, while we confirmed the expression of *TAF8s* in humans by different analyses and sample origins, we could not find evidence for *TAF8s* expression in other species, including closely related non-human primates. We conclude that *TAF8s* is very likely a *TAF8* isoform specific to the human lineage.

### *Alu* element insertion preceded iPAS formation in the human lineage

Having established that the *TAF8s* isoform is generated by the usage of an iPAS in intron 5, we further analysed this region to define the iPAS evolutionary origin. Human *TAF8* intron 5 is almost entirely constituted of transposable elements (TEs), mainly belonging to the SINE subclass (**Fig. 4A**).

**Figure 4.**
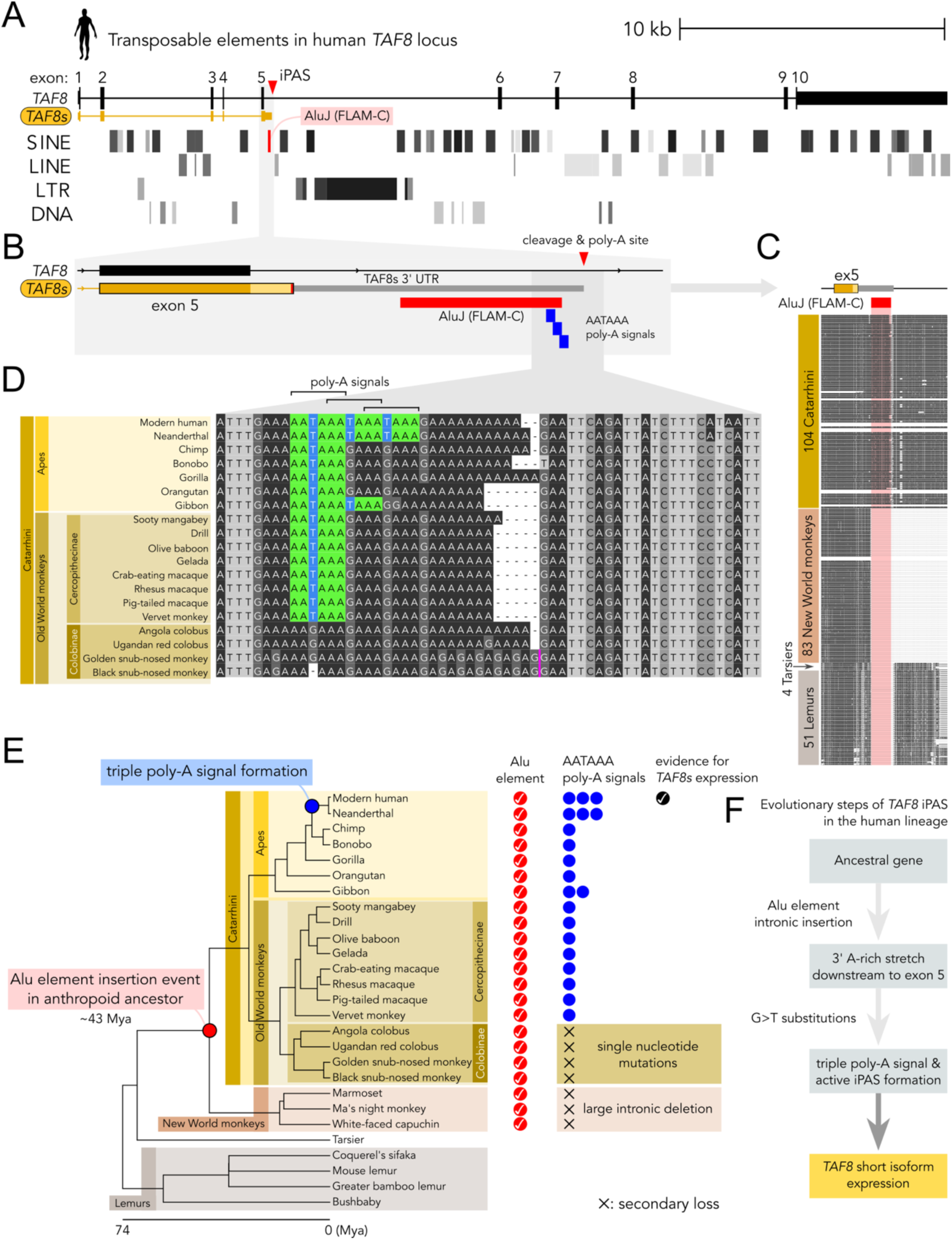
Evolutionary origin of human *TAF8* iPAS is connected to the insertion of an *Alu*. **A**. Distribution of transposable elements in the human *TAF8* locus mapped by RepeatMasker. **B**. Zoomed-in view of the genomic region encompassing *TAF8* exon 5 and downstream iPAS harbouring the three poly-A signals (in blue) in correspondence of a FLAM-C *Alu* element (in red). **C**. Multi-species primate genomes alignment track from Cactus 447-way alignment (242 species from Zoonomia genomes) surrounding *TAF8* exon 5. The position of the FLAM-C element is highlighted in red. **D**. Multiple sequence alignment of the DNA region encompassing the human *TAF8* iPAS in multiple primate species (Catarrhini). The AATAAA poly-A signals are highlighted in colour. **E**. Distribution of *Alu* elements orthologous to the human FLAM-C harbouring the iPAS across primate species. Presence of the *Alu* element is indicated by a red check mark. Each instance of a poly-A signal is indicated by a blue circle. The species are ordered according to the species tree. The inferred phylogenetic position of the *Alu* insertion event and the formation of the triple poly-A signal are indicated on the species tree. **F**. Schematic summary of the inferred evolutionary steps in the generation of the iPAS in the human lineage.

We noticed that the iPAS responsible for the generation of *TAF8s* overlaps with the very first of these SINE elements within intron 5. For instance, the three partially overlapping polyadenylation signals (AATAAA) are positioned at the end of a SINE belonging to the primate-specific *Alu* family (**Fig. 4B**). Typically, *Alu* are ∼300 bp elements consisting of two similar monomers, the left and right arms, separated by an A-rich sequence and followed by a poly-A tail (Quentin 1992). The element analysed is classified as a Free Left Alu Monomer (FLAM-C), a ∼130 bp monomeric *Alu* element of the AluJ subfamily. The alignment of the human element to the FLAM-C consensus sequence shows that the three poly-adenylation signals match to the 3’ A-rich region common to most *Alu* elements (**Fig. S6A**). This suggests that the origin of *TAF8* intronic polyadenylation signals is evolutionarily linked to the *Alu* element they are embedded in. The inspection of genomic DNA sequence alignments from 242 primates around *TAF8* exon 5 reveals that the FLAM-C element is present in the group constituted by Old World monkeys and apes (Catarrhini), while it is absent in tarsiers (Tarsiidae) and lemurs (Strepsirrhini) (**Fig. 4C**). In New World monkeys (Platyrrhini) the scenario is more complex: (i) the FLAM-C element is not found, and it is replaced by an AluSx4 element in the same position; (ii) intron 5 underwent a large internal deletion in correspondence to the above FLAM-C element. Indeed, the human-marmoset (New World monkey) genome alignment shows a 7.4 kb deletion in marmoset intron 5, with the two breakpoints mapped within the FLAM-C and an AluSx element downstream in the corresponding human intron 5 (**Fig. S6B**). These observations suggest that the common ancestor of the New World monkeys likely underwent an *Alu*/*Alu* repeat-mediated deletion within *TAF8* intron 5, caused by the recombination between the FLAM-C and AluSx elements (**Fig. S6C**). Having established that Old World monkeys and apes share the same FLAM-C element, we aligned DNA sequences from representative species to define the occurrence of the poly-A signals that drive the expression of *TAF8s* in humans (**Fig. 4D**). We also mapped Neanderthal DNA reads to the human genome and included the derived sequence in the alignment as the closest relative to our species (**Fig. S6D**). Interestingly, modern human and Neanderthal are the only species with three partially overlapping poly-A signals (AATAAA). Other apes have only one poly-A consensus sequence, except for gibbons, with two occurrences. Old World monkeys also have only one AATAAA site, except for leaf-eating monkeys (Colobinae), as this clade presents a substitution or deletion of the central T in the consensus. The analysed region also varies in the length of the poly-A stretch downstream of the poly-A signal, hinting at an inherent evolvability of this homopolymeric tract (**Fig. 4D**). We combined the above observations with the phylogenetic relationships among the species to infer the most likely sequence of events that shaped the iPAS along the human lineage (**Fig. 4E**). In the common ancestor of higher primates (New World monkeys, Old World monkeys and apes), an *Alu* insertion event occurred ∼120 bp downstream of *TAF8* exon 5, around ∼43 million years ago. The 3’ region of the *Alu* element brought an A-rich stretch that either contained an AATAAA site or facilitated its formation through subsequent mutations. The AATAAA site was independently lost in New World monkeys, due to the intronic *Alu*/*Alu* repeat-mediated deletion event, and in the Colobinae clade through single-nucleotide substitutions. Lastly, gibbons and the genus *Homo* independently acquired one and two additional AATAAA sites, respectively, through G>T substitution events. The acquisition of the triple poly-A site in the human lineage can be mapped to a broad time range, between the last human-chimp common ancestor (5-7 Mya) and the modern human-Neanderthal common ancestor (0.5 Mya). As suggested by the above gene expression analyses in primates, the presence of a single AATAAA signal is not sufficient to drive the expression of *TAF8s*. Interestingly, RNA secondary structure predictions of the region surrounding the poly-A signals unveil potential differences in RNA folding of the human sequence from that of other primates (**Fig. S7**). Specifically, the region enclosing the cleavage and poly-A site in humans is predicted to be part of an extended single-stranded region, while in other primates, this region might fluctuate more in a stem-loop state, potentially hindering access to the cleavage and poly-adenylation machinery in non-human primates. We conclude that the intronic insertion of an *Alu* element and subsequent formation of the three partially overlapping AATAAA signals – along with other more subtle changes affecting RNA secondary structure dynamics – shaped *TAF8* intron 5 in the human lineage, permitting the unique expression of *TAF8s* in humans (**Fig. 4F**).

### TAF8 short protein isoform lacks nuclear localisation signal and is mostly cytoplasmic

A key difference among the annotated functional regions between the canonical and short TAF8 isoforms is that TAF8s lacks the experimentally validated nuclear localisation signal (NLS) positioned at the C-terminus of the canonical polypeptide (pos. 294-307, **Fig. 5A**) (Soutoglou et al. 2005). Thus, we investigated the subcellular localization of the human specific TAF8s isoform. We transfected N-terminal FLAG-tagged versions of either the canonical TAF8 (TAF8c) of the TAF8 short (TAF8s) isoforms in HeLa cells and performed anti-FLAG immunofluorescence (**Fig. 5B**). For TAF8c, we observed the expected strong enrichment of signal in the nucleus. Instead, the TAF8s protein localised homogeneously throughout the cell volume. We performed the same experiment in a second, unrelated glioblastoma cell line (U87MG), confirming a lack of nuclear accumulation of TAF8s (**Fig. 5C**). To verify these observations on endogenous proteins, we performed cytoplasmic/nuclear cell fractionation experiments on three distinct human cell lines (U87MG, A172, HeLa). The two isoforms showed a clear difference in their repartition between the subcellular fractions in all cell lines, with TAF8c enriched in the nuclear fraction and TAF8s mostly partitioned in the cytoplasmic extracts (**Fig. 5D**). In summary, our experiments show that the two TAF8 isoforms differ in their subcellular localization and that TAF8s lacks nuclear tropism.

**Figure 5.**
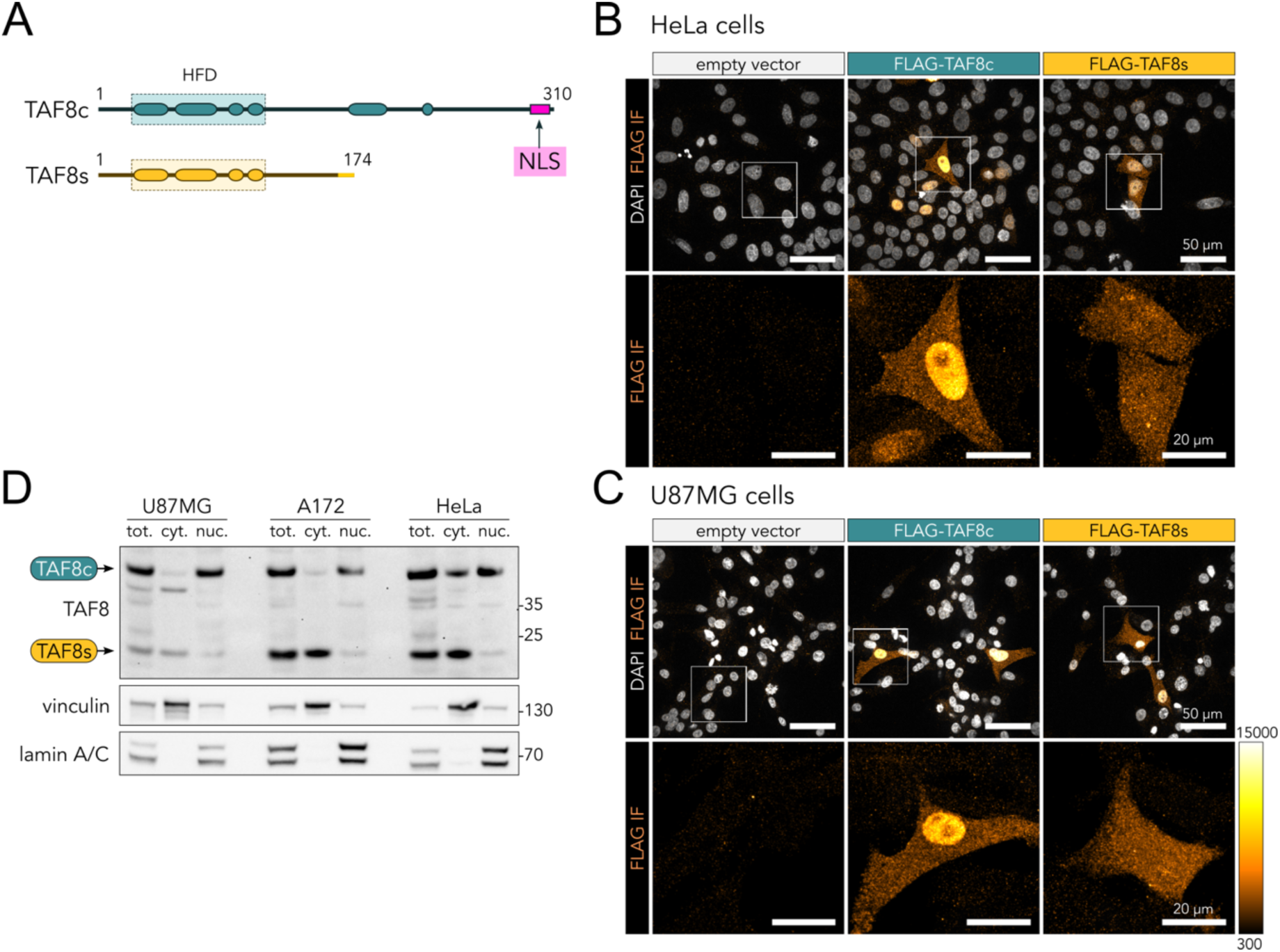
TAF8s lacks nuclear tropism. **A**. TAF8c and TAF8s protein isoforms functional domains organization. The location of the validated NLS in TAF8c isoform is highlighted in pink. **B**. Maximum intensity projections of FLAG immunofluorescence (IF, orange-hot colour scale) confocal micrographs of HeLa cells transfected with FLAG-tagged TAF8 isoforms expression vectors (as indicated). Nuclei were counterstained using DAPI (grayscale). IF signal from the square regions is enlarged in the lower panels. **C**. Same as panel B but using U87MG cells. **D**. Western blot of distinct human cell lines subjected to subcellular fractionation and probed for endogenous TAF8 isoforms. Total cellular (tot.), cytoplasmic (cyt.) and nuclear (nuc.) extracts are loaded in equal proportions. The bands corresponding to the canonical and short TAF8 isoforms are indicated. Vinculin and lamin A/C were used as cytoplasmic (cyt.) and nuclear (nuc.) markers, respectively.

### TAF8 short lacks all TAF2-interacting regions

As TFIID assembly occurs in the cytoplasm, we investigated how the TAF8 short isoform differs from the canonical isoform in terms of interactions with the other TFIID subunits. The two main TAF8 interaction partners are TAF10, with which it establishes a tight histone-fold heterodimer positioned in lobe B, and TAF2, a large globular subunit located in lobe C that participates in binding the downstream promoter DNA (**Fig. 6A**). We mapped the protein-protein interfaces along the TAF8 sequence resolved in the available experimental structures (aa 24-232). The N-terminal portion – common to both TAF8 isoforms – interacts extensively with TAF10, through the histone-fold domain (HFD), and with the core TFIID subunits TAF4-TAF5-TAF6-TAF9 in the lobe B, using the HFD (aa 24-116) and the adjacent proline-rich region (aa 117-163) (**Fig. 6B**). The C-terminal half of the protein – missing in the short isoform – mediates contacts with TAF2 (**Fig. 6B**). Specifically, TAF8c has two separate TAF2-interacting regions, of which only region 1 (aa 172-230) has been experimentally visualized. Yet, pull-downs on baculovirus-expressed complexes (Scheer et al. 2021) and tiled peptide arrays (Trowitzsch et al. 2015) identified a second TAF2-interacting region close to the very C-terminus of the protein. We conclude that TAF8s isoform lacks both TAF2-interacting regions, which were shown to contribute cooperatively to TAF2 association (Trowitzsch et al. 2015).

**Figure 6.**
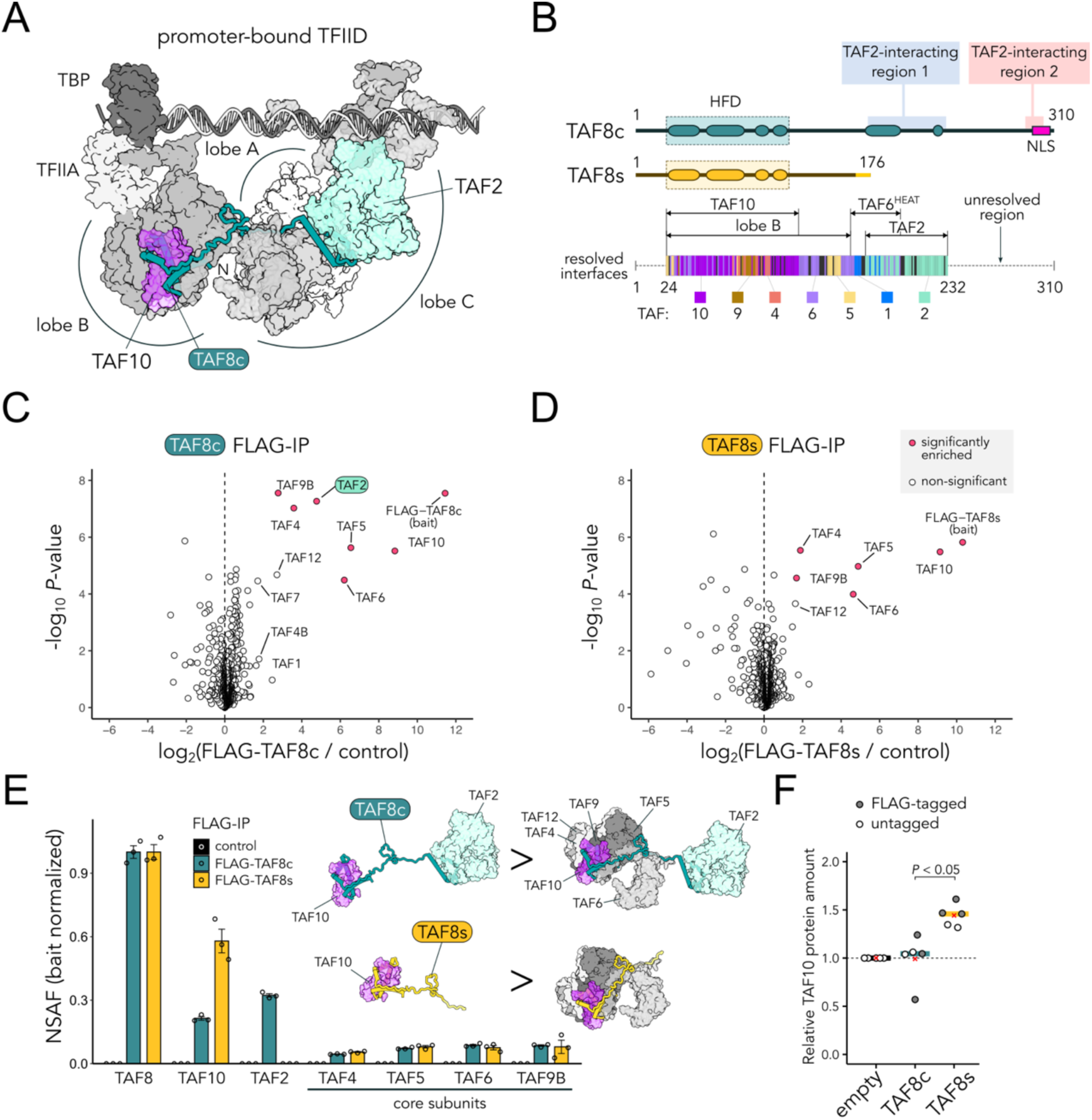
TAF8s lacks TAF2-interacting regions. **A**. Location of TAF8c protein in the DNA-bound TFIID complex structure (PDB: 7EGJ). TAF8 is depicted with ribbon representation, while other subunits are shown with surface representation. TFIID lobe A is positioned behind lobe C and coloured in white. **B**. TAF8c and TAF8s protein isoforms functional domains organization. HFD: histone-fold domain. The location of the TAF2-interacting regions 1 and 2 is indicated. TAF8 protein-protein interfaces mapped from the available experimental TFIID structures are shown in the lower panel. Each TAF8 residue is color-coded according to the main interacting subunit within the complex. Regions not resolved in the experimental structures are shown as a dotted line. **C**. Volcano plot of FLAG-IP mass-spectrometry data from HeLa whole cell extracts expressing FLAG-TAF8c. Empty vector cells were used as negative control. Significantly enriched hits are indicated. **D**. Same as panel C but for FLAG-TAF8s. **E**. Average NSAF (normalized spectral abundance factors) values of TAFs enriched in the experiments shown in panels C and D (n = 3 technical replicas). Bait NSAF values are set to one. Error bars represent SEM. The sub-complexes inferred from the estimated stoichiometry are depicted on the side. **F**. TAF10 protein quantification from western blot experiments on whole-cell extracts from HeLa cells transfected with TAF8 isoforms. TAF10 protein levels between TAF8c and TAF8s samples were compared using a paired t-test across independent experiments (n = 5), setting the empty-vector transfected sample to 1. Experiments were performed both with and without FLAG-tagged constructs (grey and white dots, respectively). Bars and red crosses correspond to the median and mean values, respectively.

### TAF8 short isoform associates with TAF10 but fails to interact with TAF2

To test the observations made from structural analyses, we carried out immunoprecipitation (IP) experiments to isolate the TFIID subunits interacting with the two TAF8 isoforms. To directly probe TAF8s interactions, we raised a polyclonal antibody (#1640 pAb) directed against the C-terminal peptide present uniquely in TAF8s isoform (**Fig. S8A**). The resulting antibody detects TAF8s in western blot assays using HeLa whole cell extracts, efficiently depletes TAF8s from the supernatant fraction of the IP, enriching both TAF8s and TAF10 in the IPs (**Fig. S8B**). Co-IPs performed on cytoplasmic extracts using the TAF8s-specific antibody and subjected to mass-spectrometric (MS) analysis retrieved TAF8s and TAF10, but not TAF2. A parallel control IP performed with an antibody directed against a common N-terminal TAF8 region (aa 14-32, 2TAU 2B8 mAb) retrieved TAF8, and both TAF10 and TAF2 (**Fig. S8C**). In our experience, TAF8 IPs might suffer from being agnostic to specific subcomplexes or even disrupt them due to steric hindrance of antibody-epitope interactions, as most of TAF8 protein is in direct contact with other subunits in TFIID (∼54% buried surface; see also **Fig. 6A**). Thus, we expressed N-terminal FLAG-tagged versions of TAF8c and TAF8s in HeLa cells, to have a common N-terminal epitope exposed to the solvent. We performed FLAG-IP on whole cell extracts (mostly representative of the cytoplasmic fraction), followed by label-free MS analysis. In TAF8c FLAG-IP, core TFIID subunits (TAF4-TAF5-TAF6-TAF9), together with TAF10 and TAF2, were significantly enriched (**Fig. 6C**). Similarly, TAF8s FLAG-IP strongly enriched TAF10, and – to a lower extent – the same core TFIID subunits. However, TAF8s entirely failed to associate with TAF2 (**Fig. 6D**), similarly to the endogenous protein IP. By analysing the abundance of the IP-enriched proteins using the normalized spectral abundance factors (NSAF) method, we infer that TAF8c forms an abundant trimeric assembly with TAF10 and TAF2, named 3TAF complex, a fraction of which is found associated with the core TFIID submodule (**Fig. 6E**). The retrieval of these TFIID building blocks recapitulates previous experiments from cytoplasmic IPs of endogenous TAFs (Trowitzsch et al. 2015; Bernardini et al. 2023). Instead, the TFIID building blocks detected in the TAF8s-specific IPs all lack TAF2 (**Fig. 6E**), consistently with the structural observations above. Moreover, in contrast to the canonical isoform, overexpression of the TAF8s isoform leads to a ∼50% increase in TAF10 protein abundance (**Fig. 6F**), suggesting a stabilization effect specific to the short isoform. This might also explain why we found higher relative levels of TAF10 complexed with TAF8s when compared to the canonical isoform IPs (**Fig. 6E**). We conclude that the newly evolved human TAF8s isoform lacks the capacity to associate with, and recruit TAF2 in canonical TFIID sub-complexes due to the absence of the specific C-terminal anchor points.

## Discussion

### The origins of a novel human-specific TAF8 pan-isoform

Here, we provide the first description of a human-specific protein isoform of a TFIID subunit. Indeed, we found no evidence for the expression of *TAF8s* in all the tested non-human species. On the contrary, *TAF8s* is ubiquitously expressed in human tissues and cell lines, where a fraction of the nascent *TAF8* transcripts is constitutively cleaved at the intronic site (iPAS) and Pol II elongation terminates prematurely, generating the shorter *TAF8s* mRNA. Thus, we refer to *TAF8s* as a human-specific pan-isoform co-expressed along with canonical *TAF8*. Nevertheless, the ratio of the two isoforms does vary in different tissues (**Fig. S1C**), opening the question of whether *TAF8s* expression is actively regulated, especially during embryonic development, or dysregulated in certain diseases (Grzechnik and Mischo 2025). As *TAF8s* is generated by an intronic cleavage and poly-adenylation event, we focused on the iPAS region of *TAF8* intron 5 to investigate the evolutionary origins of the short isoform. The three human poly-A signals are located at the 3’ extremity of a FLAM-C retrotransposon, a monomeric member of the AluJ family. By comparing DNA sequences across primate genomes, we mapped the origin of the FLAM-C insertion at the base of the anthropoid lineage (Simiiformes, which include New World monkeys, Old World monkeys, and apes), which occurred after the divergence of tarsiers from the haplorrhine stem. This timing aligns with reconstructions showing high FLAM activity in the common ancestor of anthropoids, followed by a marked reduction in retrotransposition thereafter (Schmitz et al. 2016). The FLAM-C insertion provided an A-rich sequence background, probably already containing a first AATAAA signal. The cryptic poly-A signal was independently disrupted in New World monkeys by a large intronic deletion, and in Colobinae by single-nucleotide substitutions (**Fig. 4E**), suggesting that the newly inserted transposon likely did not provide any selective advantage. We infer that the PAS remained dormant in a non-functional state for several million years, until a series of DNA changes activated it in our lineage, after the divergence from other apes. The prominent change was probably the generation of a triple partially overlapping poly-A signal that is unique to humans, including Neanderthals. Yet, it is likely that other changes in the surrounding sequence enhanced the usage of the iPAS as well, possibly through changes in the secondary structure dynamics of the nascent transcript that favoured its recognition by the cleavage and polyadenylation machinery. Additionally, changes in the splicing efficiency (Yang et al. 2025) or chromatin configuration of the locus (Spies et al. 2009) might have contributed to activating the *TAF8* iPAS in our lineage, aspects not investigated here. The case we describe is an example of the exaptation of an *Alu* element as a polyadenylation site (Lee et al. 2008; Chen et al. 2009), recently described also for non-primate SINEs in dogs (Choi et al. 2025). The case described here impacts on a subunit of a fundamental component of the core transcription machinery, adding to the multiple ways TEs can shape gene expression and evolution.

### Potential TAF8s molecular functions and connection to disease

We demonstrate here that the shorter transcript that originates from the usage of the iPAS is translated into a C-terminally truncated protein isoform. This raises the question as to what the TAF8s molecular function could be.

First, our experiments demonstrate a sharp difference in TAF8s subcellular localization when compared to the canonical isoform, with the latter being mostly nuclear (**Fig. 5**). Instead, TAF8s lacks nuclear tropism, consistently with the lack of the experimentally validated NLS (Soutoglou et al. 2005; Trowitzsch et al. 2015). Consequently, most of the short isoform localises in the cytoplasm, where it heterodimerises with its HFD partner TAF10. Indeed, TAF8s overexpression resulted in increased TAF10 protein levels, hinting at an involvement of the short isoform in regulating the cellular levels of TAF10. A fraction of the TAF8/TAF10 heterodimer associates with core TFIID subunits in larger assemblies, as it happens for the canonical isoform. Indeed, specific TFIID subcomplexes are found in the cytoplasm of human cells, as they represent intermediates in the co-translational assembly of the holo-complex on the nascent TAF1 subunit (Trowitzsch et al. 2015; Bernardini et al. 2023).

Second, we demonstrate that TAF8s does not interact with TAF2, as it lacks all the mapped TAF2-interacting regions located in the C-terminal half of the canonical protein. Thus, TAF8s does not form any TFIID submodule containing the TAF2 subunit. TAF8 represents the major anchor point for TAF2 in the available TFIID cryo-EM structures (Patel et al. 2018; Chen et al. 2021), as also shown through biochemical means (Scheer et al. 2021). A likely consequence is that TAF8s might work as a “poisoning” or “sequestering” isoform, whereby its entry into the assembly pathway leads to a dead-end TFIID complex, due to (i) the association with TAF10 and core TFIID subunits in defective multi-TAF complexes that remain stuck in the cytoplasm and (ii) the inability to recruit TAF2 in the complex.

Nevertheless, these apparently detrimental TAF8s properties might also have interesting consequences. First, TAF8s incorporation in the assembly pathway would result in sub-stoichiometric levels of TAF2 in the TFIID complex pool. In the literature, reports that sustain this prediction in human cells exist (Kaufmann et al. 1996; Cheng et al. 2024), including TAF2-lacking TFIID complexes (Kaufmann et al. 1998). Concurrently, TAF8s incorporation would lead to a substantial fraction of free TAF2, not associated with TFIID. Indeed, unlike other TAFs, TAF2 has been recently found associated with the nuclear speckles, suggesting it can work independently in specialized subnuclear compartments (Bhuiyan et al. 2025). Moreover, endogenous TAF2 used as bait in IP-MS experiments is found in excess relative to the other TFIID subunits in human nuclear extracts, suggesting that a fraction of TAF2 exists in isolation from the holo-complex (Trowitzsch et al. 2015; Bernardini et al. 2023). Thus, TAF8s might work alongside TAF8c as a pan-isoform that opens collateral paths in TFIID assembly, potentially leading to novel partial complexes that coexist with the canonical one. This molecular “splitter” function of pan-isoforms was recently proposed for an isoform of SUZ12 – a PRC2 subunit (Arecco et al. 2024). It remains to be determined if – and how – TAF2, and the partial TFIID complexes containing TAF8s, can get be imported to and potentially function in the nucleus, possibly under specific circumstances or cell states.

The discovery of TAF8s isoform has a potential impact on the interpretation of genetic disorders driven by *TAF8* mutations. For example, in the first case reported, a homozygous *TAF8* mutation (NM_138572.2: c.781–1G > A) hitting the canonical exon 8 acceptor splice site induces a frameshift in the resulting transcript (El-Saafin et al. 2018). The mutant protein was extremely unstable, and it was not detected in the patient’s fibroblasts, suggesting that the mutation acted as a gene loss-of-function. This was a stunning observation, as one would question how an individual could even develop in the absence of TAF8, given that KO mice die at the embryonic phase (Voss et al. 2000). In this regard, the present study might provide an alternative interpretation. The above mutation would not affect *TAF8s* isoform expression, as it acts downstream of exon 5, at the junction between canonical exons 7 and 8. Therefore, we foresee that the patient’s cells would still express the TAF8s protein, potentially compensating for the lack of the canonical isoform in unexpected ways.

In conclusion, our study revealed the existence of a human-specific pan-isoform of a key TFIID scaffold subunit. The lack of nuclear tropism and inability to interact with TAF2, coupled with the capacity to efficiently interact with TAF10 and other core subunits, suggests that TAF8s might work as a molecular splitter that contributes to regulate the amount of functional TFIID in human cells. Future studies will be needed to further detail the functional regulation, and potential adaptive roles to cellular fitness of TAF8s in human cells or in disease conditions.

## Materials and Methods

### Expression constructs

The coding sequence (CDS) of *TAF8s* isoform, including the N-terminal FLAG tag, was obtained by gene synthesis (Eurofins Genomics). The CDS of *TAF8c* isoform was obtained from the pXJ41-TAF8-Nter-3FLAG construct (Kamenova et al. 2019). *TAF8c* and *TAF8s* CDSs were amplified by PCR using Pfu DNA polymerase (Agilent) and sets of primers to either include or exclude the N-terminal FLAG-tag sequence. The amplicons were purified and digested with HindIII and XhoI restriction enzymes. After enzyme inactivation and purification, the digested amplicons were ligated with pcDNA3.1 expression vector using T4 DNA ligase (NEB). All constructs were verified by Sanger sequencing.

### Cell culture and transfection

All cell lines were obtained from the IGBMC cell culture platform. HeLa and HEK293T cells (ATCC) were cultured in DMEM (4.5 g L^−1^ glucose) supplemented with 10% foetal bovine serum (FBS). A172 cells (CRL-1620; ATCC) were cultured in DMEM (4.5 g L^−1^ glucose) supplemented with 10% FBS and 40 µg/mL gentamycin. U87MG cells (HTB-14; ATCC) were cultured in MEM supplemented with 10% FBS, non-essential amino acids, 1 mM sodium pyruvate and 40 µg/mL gentamycin. GM12878 cells (Coriell Cell Repositories, NIGMS, Camden, USA) were cultured in RPMI supplemented with 10% FBS and 40 µg/mL gentamycin. COS-7 cells (ATCC) were cultured in DMEM (1 g L^−1^ glucose) supplemented with 10% FBS and 40 µg/mL gentamycin. All cells were grown at 37 °C in a humidified 5% CO_2_ incubator. Cells were transfected using Lipofectamine 2000 transfection reagent (Invitrogen) diluted in Opti-MEM (Gibco). DNA/Lipofectamine complexes were incubated at room temperature for 15 min and added to the cells. Growth medium was replaced five hours after transfection.

### Cells extracts and subcellular fractionation

RIPA extracts for western blot analysis were prepared by adding RIPA lysis buffer [10 mM Tris-HCl pH 8.0, 140 mM NaCl, 1 mM EDTA, 0.1% SDS, 0.1% sodium-deoxycholate, 1% Triton X-100, 1 mM DTT, 1× protease inhibitor cocktail (PIC)] to the cell layer and scraping the cells. After 15 min on ice, the extracts were centrifuged at 12,000 ×g for 10 min at 4°C and the supernatant was quantified using Bradford protein assay (Biorad). To prepare whole cell extracts (WCE) for immunoprecipitation (IP) experiments, cells were detached using trypsin and collected by centrifugation in complete medium. Cell pellet was washed using cold PBS and resuspended in one packed-cell-volume (PCV) of cold extraction buffer (20 mM Tris-HCl pH 8.0, 20% glycerol, 0.4 M KCl, 1 mM DTT, 1× PIC). The cell suspension was snap-frozen in liquid nitrogen and thawed on ice for three times. The extract was diluted using 4.5 PCV of IP100 buffer (25 mM Tris-HCl pH 8.0, 10% glycerol, 100 mM KCl, 5 mM MgCl_2_, 0.1% NP-40, 1 mM DTT, 1× PIC) and centrifuged at 12,000 ×g for 10 min at 4°C. The supernatant quantified using Bradford protein assay constitutes the WCE used for FLAG-IPs. To prepare cytoplasmic extracts used for endogenous IP experiments, cells were resuspended in 4 PCV of 50 mM Tris-HCl pH 8.0, 1 mM EDTA, 0.1% NP-40, 1 mM DTT, 1× PIC. After 30 min on ice, cells were lysed with 10 hits in a Dounce homogenizer and centrifuged for 10 min at 2,700 ×g at 4°C. The supernatant quantified using Bradford protein assay constitutes the cytoplasmic extract used in IPs. To performed subcellular fractionation experiments, cells were detached using trypsin and collected by centrifugation in complete medium. Cells were resuspended in cold PBS and split into two aliquots. One aliquot was lysed in 2× SDS sample buffer (200 mM DTT, 100 mM Tris-HCl pH 6.8, 4% SDS, 20% glycerol, 0.2% bromophenol blue) to obtain the total cell extract. The second aliquot was resuspended in four PCV of hypotonic buffer (10 mM HEPES pH 8.0, 10 mM KCl, 1.5 mM MgCl_2_, 0.1% Triton X-100, 1× PIC) and kept on ice for 5 min. Cytoplasmic fraction and nuclear pellet were obtained by centrifugation at 2,300 ×g at 4°C for 5 min. The nuclear pellet was resuspended in 2× SDS sample buffer. To reduce viscosity before gel loading, samples were briefly sonicated. Equal proportions of each fraction were loaded for western blot analysis.

### Immunoprecipitation coupled to mass spectrometry (IP-MS)

For FLAG-IP experiments, HeLa cells were harvested 48 hours after transfection with FLAG-TAF8 expression constructs or empty plasmid as control. Whole cell extracts (WCE) were precleared using protein-G Sepharose beads in rotation for 30 min at 4°C. Precleared extracts (0.5 mg per IP) were incubated with 10 µL ChromoTek DYKDDDDK Fab-Trap Agarose beads (Proteintech) in rotation for 2 hours at 4°C. Beads were washed twice with 0.5 mL IP100 buffer for 5 min each at 4°C and twice with IP100 buffer devoid of NP-40. Beads were transferred to new tubes and washed twice in PBS and processed for on-beads digestion for mass-spectrometry analysis: the proteins retained on the beads were reduced 30 min at 56°C with 5 mM TCEP in Urea 2M, 0.1 M Tris pH 8.0, then alkylated 30 min at room temperature with 10 mM iodoacetamide. Trypsin digestion was carried out at 37°C, overnight. For endogenous IP experiments, HeLa cytoplasmic extracts (3 mg per IP) or WCE (1.25 mg per IP) were precleared using a 1:1 mix protein-A/protein-G Sepharose beads in rotation for 30 min at 4°C. Precleared extracts were incubated in rotation for 4 hours at 4°C with 16 µL protein-A/protein-G Sepharose beads mix pre-coupled with 20 µg of the following antibodies: TAF8 (2TAU 2B8), TAF8s (#1640), rabbit IgG (mock). Beads were washed twice with 0.5 mL IP500 buffer (IP100 buffer containing 500 mM KCl) for 5 min each at 4°C and twice with IP100 buffer. Beads were transferred to new tubes and proteins were eluted in acidic conditions using 0.1 M glycine pH 2.8 and neutralized using 1 M Tris-HCl pH 8.0. Protein mixtures were TCA-precipitated overnight at 4°C, pellets were washed twice with 1 mL cold acetone, dried and dissolved in 2 M urea in 0.1 mM Tris-HCl pH 8.5 for reduction (5 mM TCEP, 30 min) and alkylation (10 mM iodoacetamide, 30 min). Trypsin digestion was carried out at 37°C overnight. Extracted peptides were then analysed using an Ultimate 3000 nano-RSLC (Thermo Scientific) coupled in line with an Orbitrap Exploris 480 via a nano-electrospray ionization source (Thermo Scientific, San Jose California) and the FAIMS pro interface. Peptides were separated on a C18 Acclaim PepMap nano-column (75 µm ID x 25 cm, 2.6 µm, 150Å, Thermo Fisher Scientific) with a 23 minutes linear gradient from 7% to 35% buffer B (A: 0.1% FA in H_2_O; B: 0.1% FA in 80% ACN, 400 nL/min, 50°C) followed by a regeneration step at 90% B and an equilibration at 7% B. The total chromatography was 30 minutes. The mass spectrometer was operated in positive ionization mode in Data-Dependent Acquisition (DDA) with FAIMS compensation voltages set to CV = -45V. The DDA cycle consisted of one survey scans or MS1 (300-1500 m/z, 120,000 FWHM) followed by MS² spectra (HCD; 30% normalized energy; 10 m/z window; 30,000 FWMH) in the limit of 1.2 sec. The Automatic Gain Control (AGC target) were set to 3E6 and 1E5 respectively for MS1 and MS2 scans, and the maximum injection time (IT) was set to 50 ms for both scan modes. Unassigned and single charged states were rejected. The exclusion duration was set for 40 s with mass width was ± 10 ppm.

### Mass spectrometry data analysis

Proteins were identified with Proteome Discoverer 2.5 software (Thermo Fisher Scientific) and *Homo sapiens* proteome database (UniProt, reviewed, release 2022_10_27 with 20607 sequences). Precursor and fragment mass tolerances were set at 10 ppm and 0.02 Da respectively, and up to 2 missed cleavages were allowed. Oxidation (M) and N-terminal Acetylation were set as variable modification, and Carbamidomethylation (C) as fixed modification. Peptides were filtered with a false discovery rate (FDR) at 1%, rank 1. Proteins were quantified with a minimum of 1 unique peptide based on the XIC (sum of the Extracted Ion Chromatogram). The quantification values were exported in Perseus (2.0.11.0) for statistical analysis involving a log_2_ transform, normalization on the median, imputation of missing values and followed by t-test calculation (FDR=0.01). Normalized spectral abundance factors (NSAF) were calculated as previously described (Zybailov et al. 2006).

Briefly, to obtain spectral abundance factors (SAF), spectral counts identifying a protein were divided by the protein length in amino acids. To calculate NSAF values, the SAF values of each protein were divided by the sum of SAF values from all proteins detected in each run.

### Western blot

Samples were loaded either on home-made SDS–PAGE gels run in Tris-glycine buffer or on precast NuPAGE Bis-Tris 4-12% gradient gels run in MES buffer. Proteins were transferred on nitrocellulose membrane and probed with the following primary antibodies: TAF8 (HPA031731, Sigma-Aldrich), TAF8s (#1640, this study), TAF1 (ab264327, Abcam), TAF6 (25TA 2G7, IGBMC), TAF10 (6TA 2B11, IGBMC), Lamin A/C (sc-7292, Santa Cruz Biotechnology) and Vinculin (V284, Sigma-Aldrich). To reprobe the membrane with an antibody raised in a different species, the previous secondary antibody was inactivated with 10% acetic acid according to (Han et al. 2020). HRP-conjugated secondary antibody chemiluminescence was detected using a ChemiDoc Touch system (BioRad) and images were visualized and processed in ImageLab software (BioRad).

### Immunofluorescence

Transfected cells seeded on coverslips (no. 1.5H, Marienfeld) in a 12-well plate were washed twice with PBS and fixed with 4% paraformaldehyde (Electron Microscopy Sciences) in PBS for 10 min at RT. Cells were washed twice with PBS and incubated for 10 min at RT in blocking/permeabilization solution (BPS) (1× PBS, 1% BSA, 0.1% Triton-X100). Cells were incubated for 2 hours at RT with the anti-FLAG antibody (DYKDDDDK tag mAb, Proteintech 66008-4-Ig) diluted 1:1000 in BPS. After three 5-min PBS washes, samples were incubated for 1 hour at RT with AF488-conjugated secondary antibody diluted 1:3,000 in BPS (goat anti-mouse IgG, A11001, Life Technologies) including 0.5 μg/mL DAPI (Merck, MBD0015) for nuclear counterstain. After three 5-min PBS washes, coverslips were mounted with 5 μL Vectashield (Vector Laboratories, H-1000) and sealed with nail polish. Cells were imaged using spinning disk confocal microscopy on an inverted Leica DMi8 equipped with a CSU-W1 confocal scanner unit (Yokogawa), with a 1.4-NA ×63 oil-objective (HCX PL APO lambda blue) and an ORCA-Flash4.0 camera (Hamamatsu). DAPI and AF488 (IF) were excited using a 405 nm and a 488 nm laser line, respectively. Three-dimensional image acquisition was managed using MetaMorph software (Molecular Devices). Images of 2,048 × 2,048 pixels (16-bit) were acquired with a xy pixel size of 0.103 μm and a z step size of 0.3 μm. Maximum intensity projections (MIPs) and were generated in Fiji (Schindelin et al. 2012) and image panels were generated using FigureJ plugin (Mutterer and Zinck 2013).

### Polyclonal TAF8s antibody production and purification

The peptide corresponding to the TAF8s C-terminal tail (aa 161-174) and including a N-terminal Cys residue (CKTPVSDEALGLRVV) was produced with solid phase synthesis (IGBMC peptide synthesis service). The peptide was coupled to maleimide-activated ovalbumin (Thermo Scientific) and it was used as antigen for custom rabbit immunization service for polyclonal antibody production (Agro-Bio). For antibody purification, the peptide used as antigen was coupled to SulfoLink Coupling Resin (Thermo Scientific) and post-immune rabbit serum was incubated with the resin-coupled peptide on a chromatography column. After PBS washes, the polyclonal antibody was eluted in 0.1 M glycine pH 2.8 and neutralized using 1.5 M Tris-HCl pH 8.8. The resulting fractions were pooled, dialyzed in PBS and tested in western blot and IP experiments. The obtained polyclonal antibody is named TAF8s #1640.

### Protein-protein interface analysis and structural visualization

Residue-level protein-protein interfaces from experimental structures of TAF8 were downloaded from PDBe-KB (https://www.ebi.ac.uk/pdbe/pdbe-kb/) and were mapped on the primary structure using the drawProteins R package (Brennan 2018). All structural models were visualized and rendered in UCSF ChimeraX (Pettersen et al. 2021).

### Sequence alignments, gene tracks and RNA secondary structure prediction

Multiple sequence alignments (MSA) were generated and edited using Jalview (Waterhouse et al. 2009). Gene tracks and RNA-seq tracks were visualized in the UCSC Genome Browser (https://genome.ucsc.edu/) or in IGV (Robinson et al. 2011). Species tree were generated using TimeTree (Kumar et al. 2017). RNA secondary structure predictions were computed using Vienna RNAfold web server v2.6.3 (Gruber et al. 2008) with default parameters, using 180-nt sequences centred on the first AAUAAA poly-A signal of human *TAF8* iPAS (hg38 chr6:42057674-42057853) and orthologous sequences from chimp, gorilla and gibbon.

### Gene expression analyses

The proportion expressed across transcripts (*pext*), described in (Cummings et al. 2020), for *TAF8* (**Fig. 1B**) was downloaded from gnomAD (https://gnomad.broadinstitute.org/downloads) and the mean *pext* value across human tissues was calculated exon-wise. Short-read RNA-seq data were first downloaded using the SRA Explorer website (https://sra-explorer.info/), then mapped with STAR (version 2.7.8a) (Dobin et al. 2013). Mapped read coverage was visualized by loading the BAM file corresponding to each aligned sample into IGV. For chimpanzee, macaque and lemur we quantified exon-exon junction coverage using the *SJ.out.tab* file provided as output by STAR. To guide the analysis, we used a modified GTF annotation that included a *TAF8s* homolog for each species. In Fig 3C, counts spanning each junction are represented as percentage of the total junction-spanning reads. We performed all the analyses downstream of the expression quantification in the R programming environment (version 4.0.3).

The full-length cDNA *isoseq* read collections from chimpanzee, bonobo, gorilla, orangutan, Siamang and Hoolock gibbon were first mapped using minimap2 (version 2.24-r1122) (Li 2018), employing the *splice* preset and the options *-secondary = no, -uf, -C5*. Resulting SAM files were sorted using samtools (version 1.10) (Danecek et al. 2021), then processed with the script *collapse_isoforms_by_sam.py* from the Cupcake collection (https://github.com/Magdoll/cDNA_Cupcake). Finally, the collapsed, unique, full transcripts were visualized with IGV.

## Data availability

The datasets presented in the article are derived from sources in the public domain. The mass spectrometry proteomics data have been deposited to the ProteomeXchange Consortium via the PRIDE partner repository (https://www.ebi.ac.uk/pride/) with the dataset identifier PXD072860 (<u>made available upon publication</u>). Human *TAF8* isoforms gene expression data shown in **Fig. 1A** are available at the GTEx portal (https://gtexportal.org/home/). 3’-seq tack shown in **Fig. 1C** is available at the PolyASite database v3.0 (https://polyasite.unibas.ch/) (Moon et al. 2025). Aggregate (Ribo-seq) signal track shown in **Fig. 2A** is available at the GWIPS-viz browser (http://gwips.ucc.ie/) (Michel et al. 2018). Polyadenylation site predictions using PolyA Detector are available at APARENT (https://apa.cs.washington.edu/detect) (Bogard et al. 2019). RNA-seq tracks shown in **Fig. 3A-B** are available at GEO (https://www.ncbi.nlm.nih.gov/geo/) with accession number GSE106580 (Li et al. 2018). Human splice-junction coverage data in **Fig 3C** were obtained from the GTEx portal (v8 release, GTEx_Analysis_2017-06-05_v8_STARv2.5.3a_junctions.gct.gz); short-read RNA-seq data used for the same analysis in Chimpanzee, Macaque and Mouse Lemur are from (Peng et al. 2015; Xu et al. 2018) and The Kunming Institute of Zoology, under accession numbers PRJNA271912, PRJNA393104 and PRJNA761017, respectively.

Long-reads RNA-seq data shown in **Fig 3D** were retrieved from (Yoo et al. 2025) under the accession PRJNA1016395, while human cell lines data in **Fig S4** were downloaded from ENCODE (Kagda et al. 2025), searching for long-read RNA-seq experiments.

Multi-species primate genome Cactus 447-way alignment shown in **Fig. 4C** is available from the UCSC Genome Browser. Organisms silhouettes were downloaded from PhyloPic (https://www.phylopic.org/).

## Acknowledgements

This work was supported by the Fondation ARC pour la recherche sur le cancer (ARCPOST-DOC2021080004113 fellowship awarded to A.B.). This work was financially supported by Agence Nationale de la Recherche (ANR) ANR-PRCI-19-CE12-0029-01 (EpiCAST); ANR-20-CE12-0017-03, ANR-22-CE11-0013-01_ACT; Fondation pour la Recherche Médicale (EQU-2021-03012631); NIH MIRA (R35GM139564); and NSF (Award Number:1933344) grants (to L.T. and S.D.V.). A.B. was affiliated with the IGBMC and the Université de Strasbourg at the initial and central phases of the investigation. We thank the Imaging Center and the Proteomics Platform of the IGBMC.

