## Supplementary Figures for "A recently evolved TAF8 isoform arising from an *Alu* insertion increases TFIID assembly complexity in the human lineage"

### Supplementary Figure S1

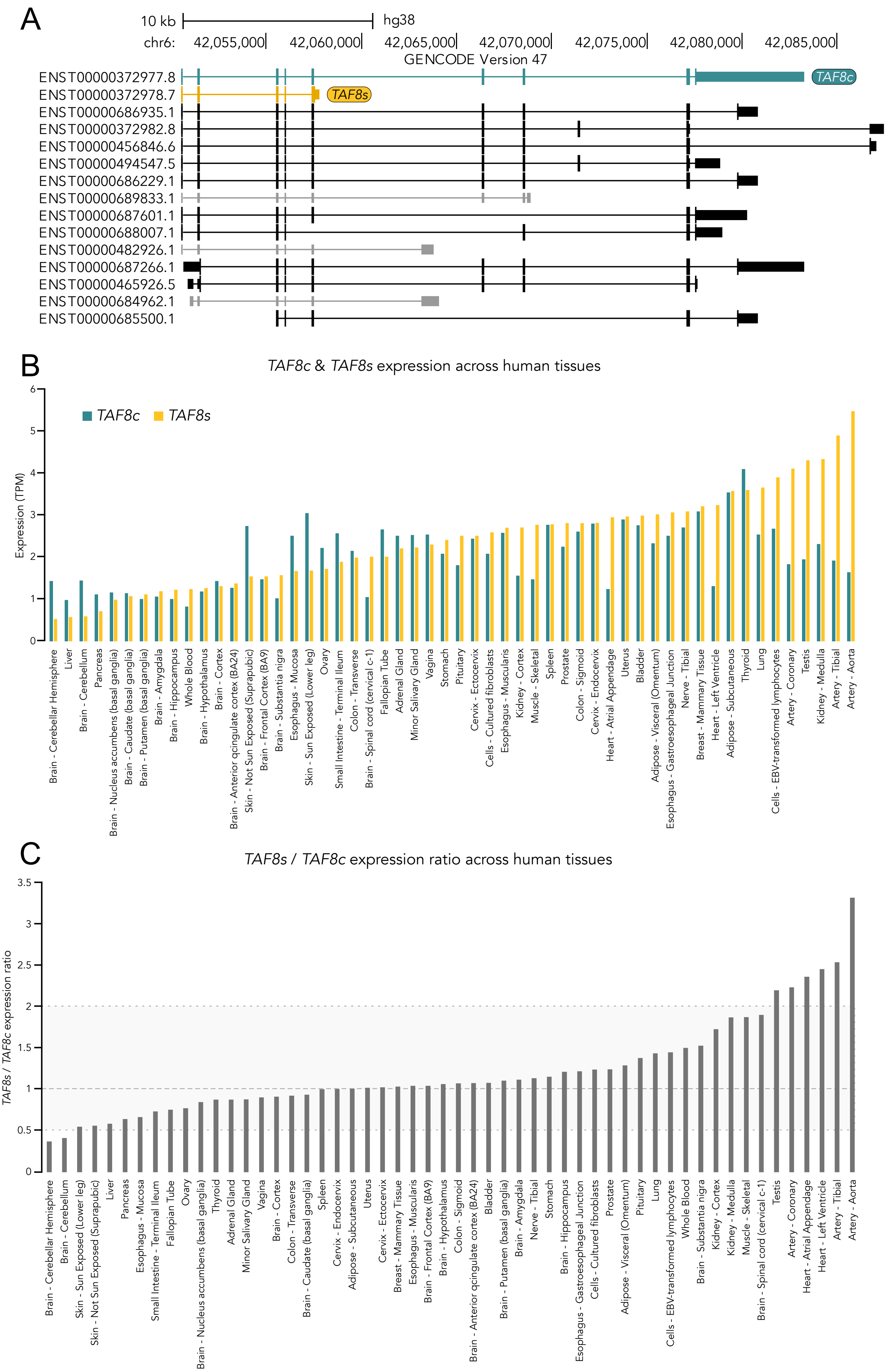

**Supplementary Figure S1. A**. Exon-intron structure of the annotated human *TAF8* alternative isoforms. *TAF8c* and *TAF8s* isoforms are highlighted in teal and yellow, respectively. Other protein-coding isoforms are shown in black and non-coding isoforms are shown in grey. Each isoform is named according to the ENSEMBL transcript ID.
**B**. *TAF8c* and *TAF8s* mRNA expression levels (expressed as transcripts per million – TPM) from human tissues of the GTEx project. Tissues are ordered according to *TAF8s* expression. **C**. *TAF8s* to *TAF8c* expression ratios calculated from data reported in panel B. Tissues are ordered according to increasing ratios.

Supplementary Figure S2

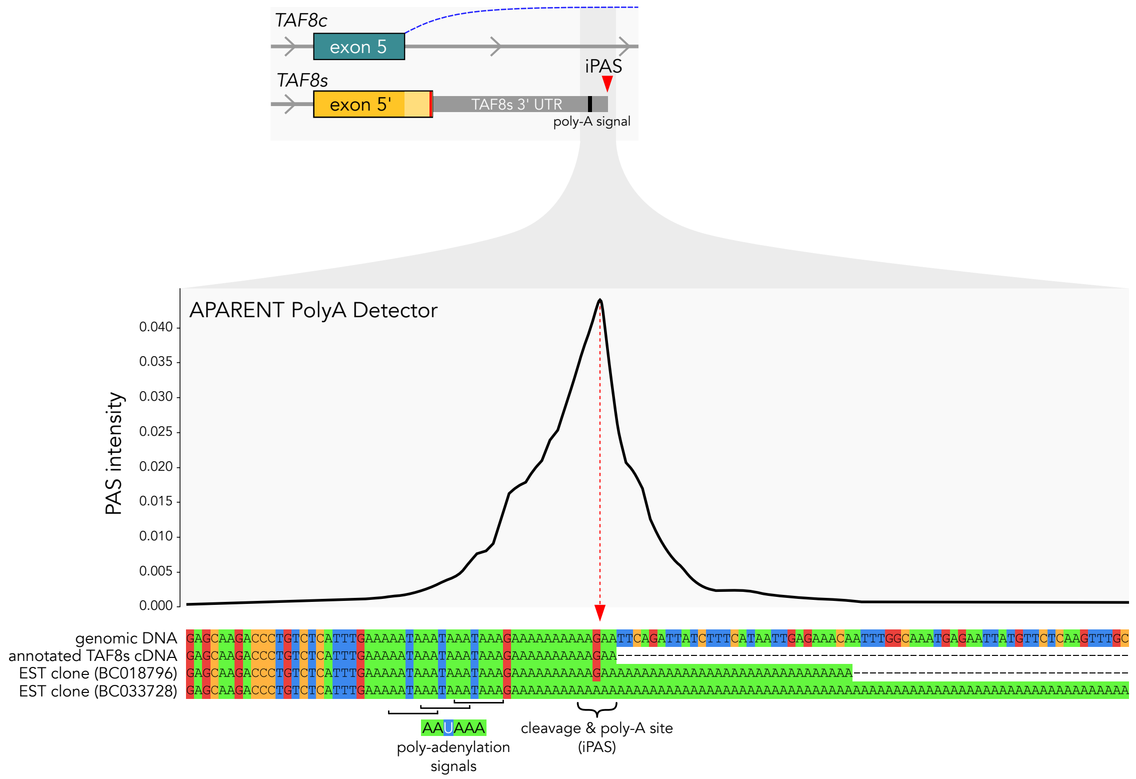


**Supplementary Figure S2.** *TAF8s* intronic polyadenylation site (iPAS) prediction by APARENT PolyA Detector (https://apa.cs.washington.edu/detect). The smoothed predicted 3' cleavage distribution is plotted along the DNA sequence around the iPAS with the significant polyadenylation peak highlighted by a vertical arrow. An alignment of the genomic DNA, the annotated *TAF8s* cDNA and two different EST clones is shown below the plot.

Supplementary Figure S3

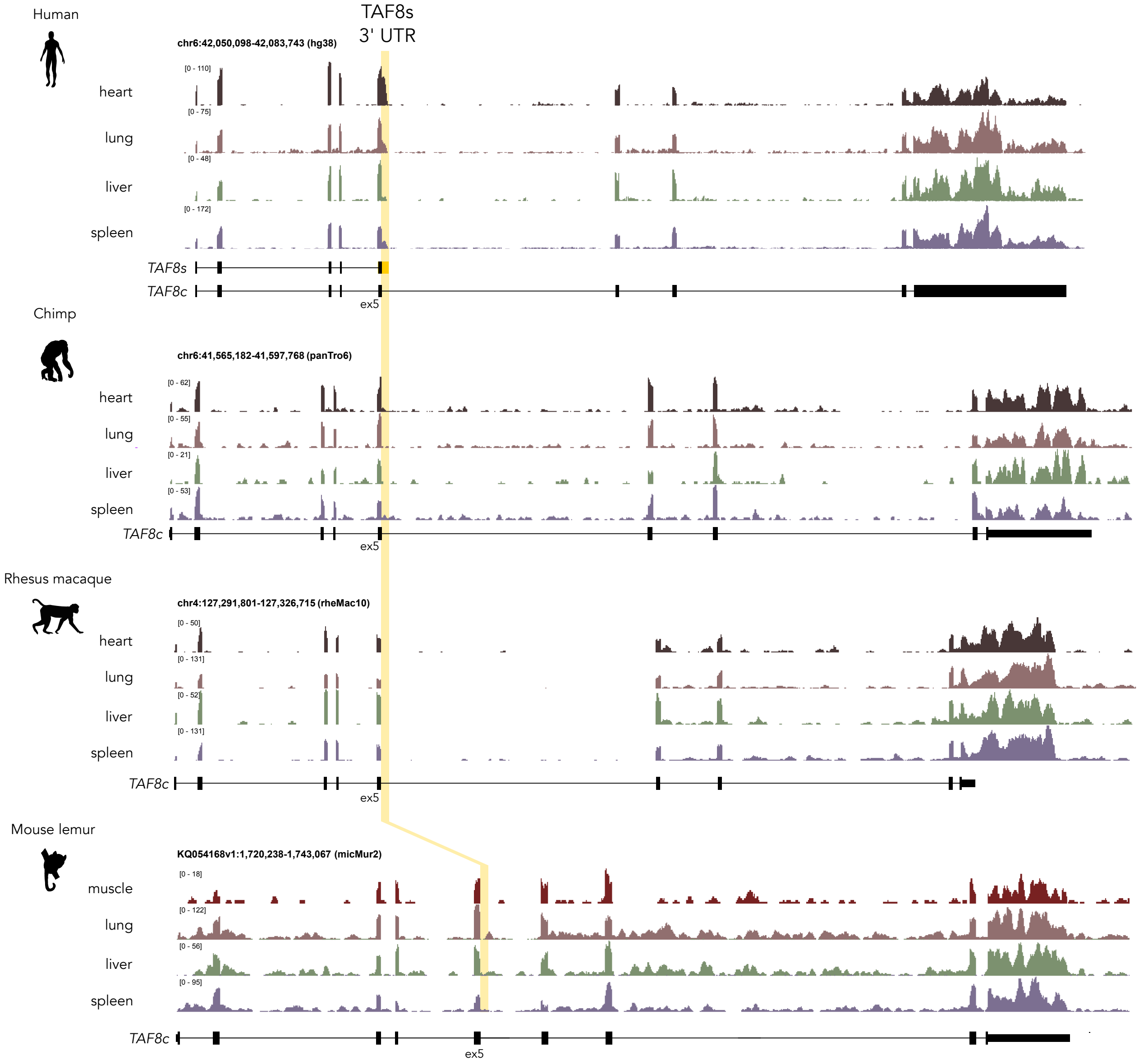


**Supplementary Figure S3.** Examples of RNA-seq coverage tracks from different primates shown in Fig. 3C. The region corresponding to the expected position of *TAF8s*-specific 3'UTR is highlighted in yellow.

Supplementary Figure S4

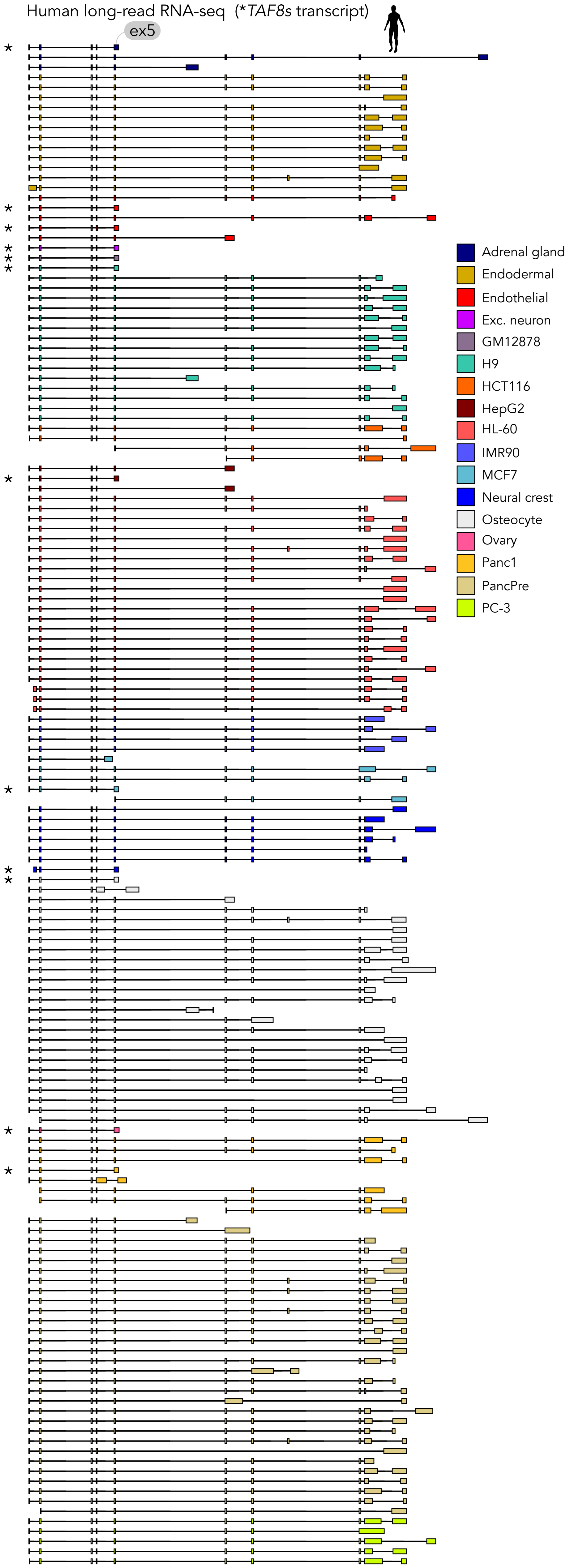


**Supplementary Figure S4.** Exon-intron structures of individual long reads mapped on *TAF8* gene from human samples processed with long-read RNA-seq data. Reads are grouped by tissue of origin. Instances of reads matching *TAF8s* isoform are indicated by asterisks.

Supplementary Figure S5

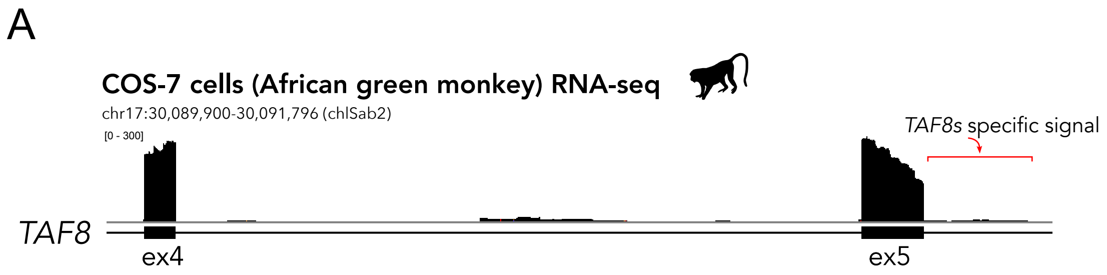


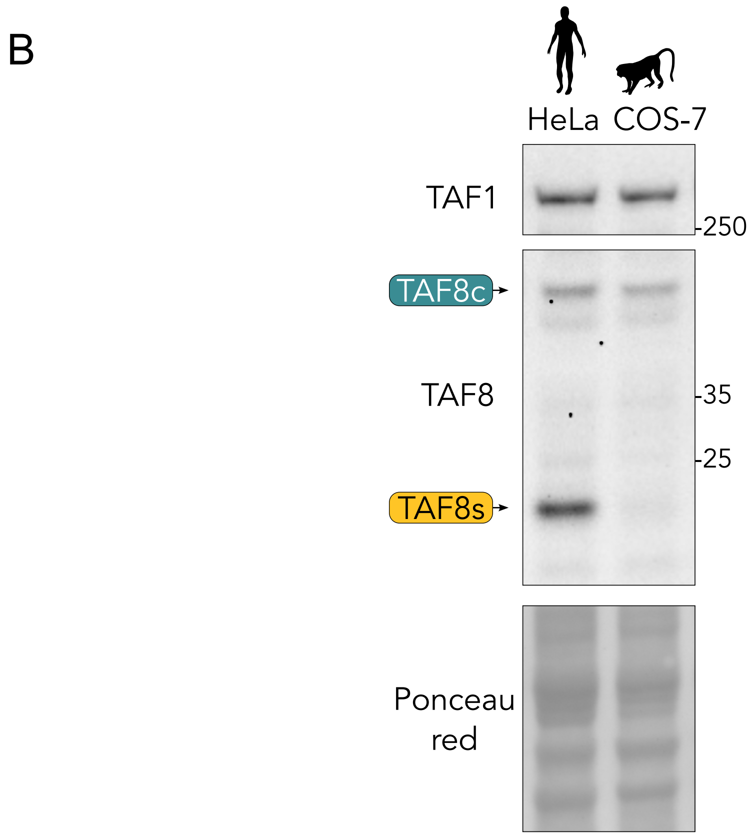


**Supplementary Figure S5. A**. RNA-seq coverage track of the African green monkey cell line COS-7. The region displayed corresponds to *TAF8* gene region encompassing exons 4 and 5. No signal is detected in the *TAF8s*-specific region. **B**. Western blot on RIPA extracts from human (HeLa) and African green monkey (COS-7) cells probed using the TAF8 antibody, showing a lack of TAF8s expression in monkey cells.

#### Supplementary Figure S6

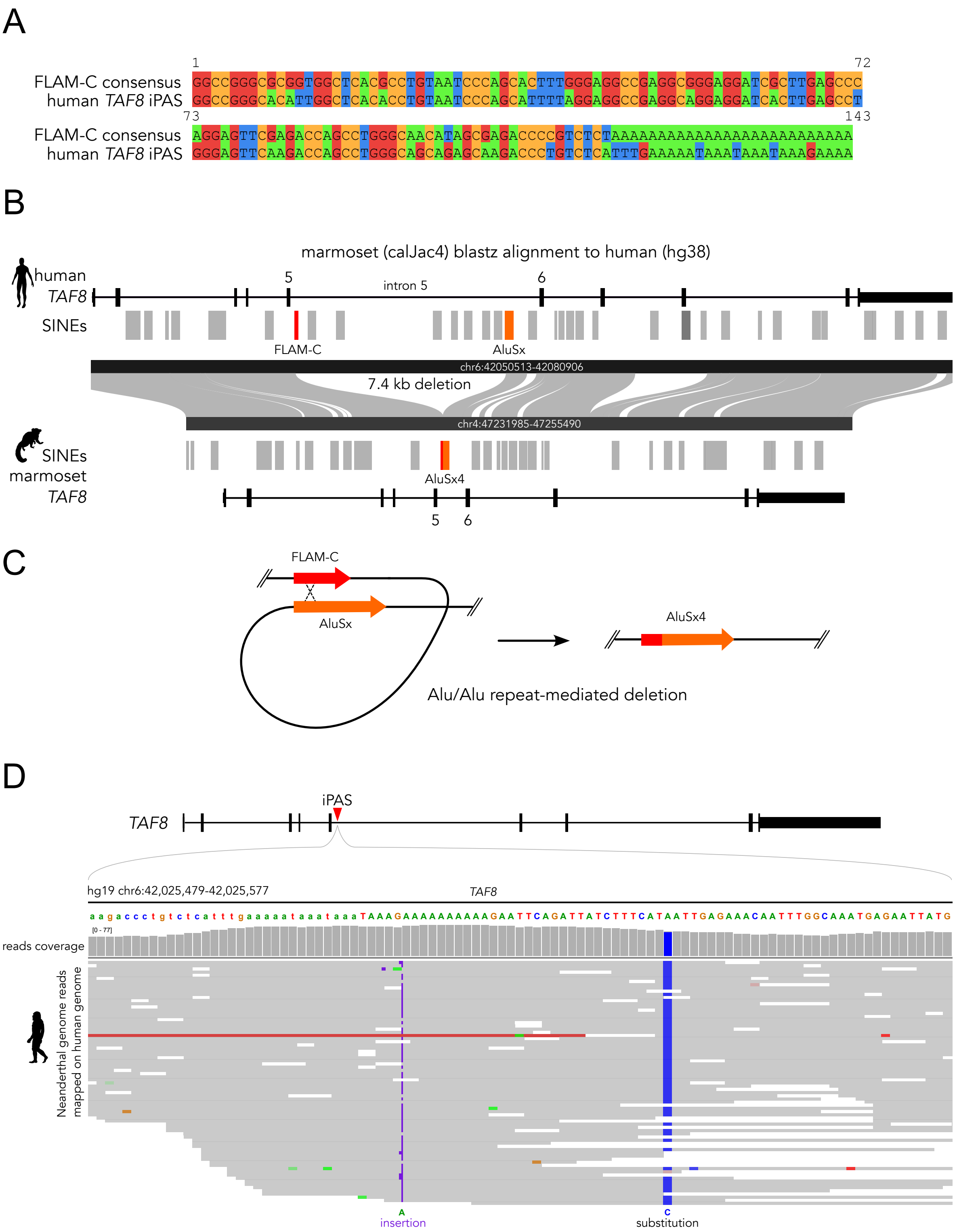


**Supplementary Figure S6. A**. Alignment of the human *Alu* element overlapping *TAF8* iPAS and the FLAM-C TE consensus sequence obtained from the Dfam database. **B**. *TAF8* locus alignment between human and marmoset genomes visualized using the WashU Comparative Epigenome Browser (https://comparativegateway.wustl.edu/). The two *Alu* elements mapped as intronic deletion breakpoints in the marmoset genome are coloured in red and orange. **C**. Scheme of an *Alu-Alu* repeat-mediated deletion event. **D**. Genome browser snapshot of Neanderthal DNA reads mapped onto *Homo sapiens* *TAF8* iPAS region. The positions differing between the two species are indicated below the piled-up reads.

Supplementary Figure S7

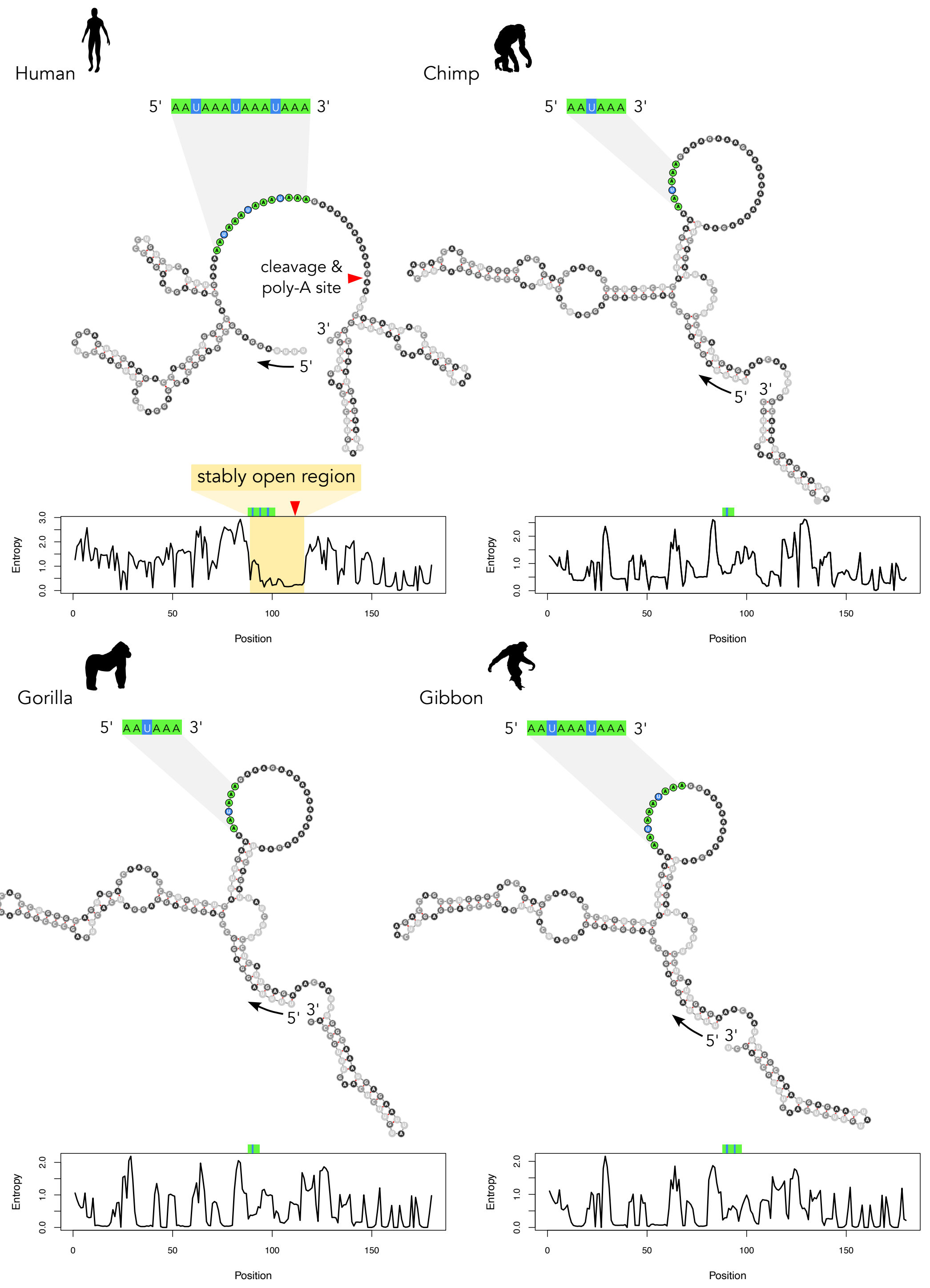


**Supplementary Figure S7.** RNA secondary structure predictions for human *TAF8* iPAS and orthologous regions of other representative primates. Predictions were obtained using the Vienna RNAfold web server (v2.6.3) with default parameters, and the minimum free energy structures are shown. A positional entropy plot is shown for each sequence, where low entropy values correspond to higher confidence in the base-pairing status. The positions of the canonical AAUAAA poly-A signals are indicated.

Supplementary Figure S8

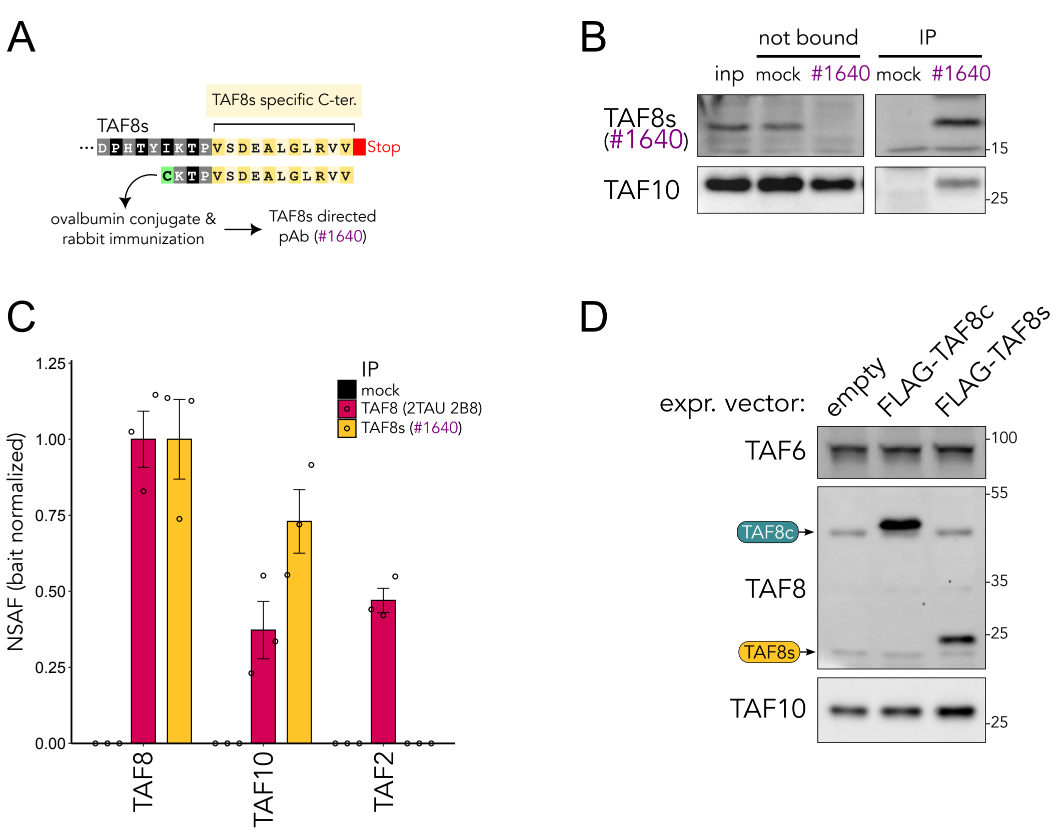


**Supplementary Figure S8. A**. Scheme of the synthetic peptide sequence matching the TAF8s-specific C-terminal tail used to generate a polyclonal antibody (pAb #1640) directed against TAF8c. **B**. Western blot of the validation of TAF8s #1640 antibody in IP with HeLa whole cell extracts. The samples include input (inp), not bound, and IP. The membrane was probed with TAF10 and TAF8s antibodies. **C**. Average NSAF (normalized spectral abundance factors) values of TAFs enriched in the IP-MS experiment on HeLa cytoplasmic extracts using TAF8s antibody (#1640) and an antibody recognizing both TAF8 isoforms (2TAU 2B8). Bait NSAF values are set to one (n = 3 technical replicas). Error bars represent SEM. **D**. Representative western blot experiment to measure TAF10 relative protein levels upon transfection of FLAG-TAF8c and FLAG-TAF8s isoforms in HeLa cells.
